# Spatiotemporal transcriptomic analysis during cold ischemic injury to the murine kidney reveals compartment-specific changes

**DOI:** 10.1101/2025.05.25.654911

**Authors:** Srujan Singh, Shishir Kumar Patel, Ryo Matsuura, Dee Velazquez, Zhaoli Sun, Sanjeev Noel, Hamid Rabb, Jean Fan

**Author notes:** Correspondence should be addressed to: Hamid Rabb, Jean Fan.

## Abstract

**Background:** Kidney transplantation is the preferred treatment strategy for end-stage kidney disease. Deceased donor kidneys usually undergo cold storage until kidney transplantation, leading to cold ischemia injury that may contribute to poor graft outcomes. However, the molecular characterization of potential mechanisms of cold ischemia injury remains incomplete.

**Results:** To bridge this knowledge gap, we leveraged the 10x Visium spatial transcriptomic technology to perform full transcriptome profiling of murine kidneys subject to varying durations of cold ischemia typical in a deceased donor kidney transplant setting. We developed a computational workflow to identify and compare spatiotemporal transcriptomic changes that accompany the injury pathophysiology in a tissue compartment-specific manner. We identified proportional enrichment of oxidative phosphorylation (OXPHOS) genes with increasing duration of cold ischemia injury within the oxygen-lean inner medulla region, suggestive of atypical metabolic presentation. This was distinct in cold ischemia injury tissue compared to warm ischemia-reperfusion kidney injury tissue. Spatiotemporal trends were validated by qPCR and immunofluorescence in a larger cohort of mice. We provide an interactive online browser at https://jef.works/CellCarto-ColdIschemia/ to facilitate exploration of our results by the broader scientific and clinical community.

**Conclusions:** Altogether, our spatiotemporal transcriptomic analysis identified coordinated molecular changes within metabolic pathways such as OXPHOS deep within the cold ischemic kidney, highlighting the need for increased attention to the inner medulla and potential opportunities for new insights beyond those available from superficial biopsy-focused tissue examinations.

## Background

Kidney transplantation remains the gold standard treatment strategy for end-stage kidney disease, relying on viable allografts from either living or deceased donors. Although live donor kidneys usually have better short and long-term graft outcomes, their limited availability has led to a major push to expand the deceased donor pool (*1*). However, a key challenge with deceased donor kidneys is cold ischemia time (CIT), the period during which the kidneys are preserved in cold storage after procurement and before transplantation (*2–4*). To minimize cellular injury and preserve the organ viability during transit from procurement to transplantation centers, kidneys are usually flushed with cryopreserving solution and kept on ice. During this period, kidneys experience cold ischemia injury (*5*), which can lead to delayed graft function and poor graft outcomes (*6*).

The pathophysiology of cold ischemia induced kidney injury is incompletely understood. While prior studies (*7*–*9*) have sought to characterize processes that are believed to accompany cold ischemia and lead to cellular injury, they usually do not account for the complex structure of kidney tissue. Moreover, these studies primarily investigated only a few targeted cold ischemia related processes and their associated biomarkers, potentially overlooking other clinically relevant pathophysiologic and adaptive events. For example, Salahudeen et al. (*9*) highlighted a few key biological insights into cold-induced cellular changes *in vitro* like cell necrosis, mitochondrial swelling, and generation of free radicals (superoxide and hydroxyl radicals). These characterizations remained mostly confined to the cells of the kidney cortex and did not capture the full spectrum of the biological changes accompanying cold ischemia injury. Additionally, findings from several large cohort, multi-variate retrospective studies (*6*, *10*–*14*) lack consensus on a standard clinically acceptable CIT for kidney allografts. Collectively, this highlights the need for a more detailed molecular characterization of cold ischemia injury of the entire kidney to elucidate the underlying biological processes that may contribute to poorer outcomes with increasing CIT.

Data-driven, high-throughput sequencing approaches can surveil the genome-wide molecular landscape during disease pathogenesis. In particular, advances in spatial transcriptomics (ST) technologies (*15–17*) now enable full transcriptome spatially resolved profiling of gene expression patterns. Application of such technologies coupled with data-driven computational exploration can facilitate the identification of potential biomarkers and therapeutic approaches to prolonged cold ischemia injury.

We therefore leveraged the 10x Visium ST technology to perform full transcriptome profiling of murine kidneys subject to varying durations of cold ischemia to characterize spatiotemporal molecular changes across typical durations of cold ischemia in a deceased donor kidney transplant setting (Fig 1A). Our integrative spatiotemporal computational analyses delineated trends that facilitated the identification of differential spatiotemporal molecular dynamics in a compartment-specific manner to shed light on how prolonged cold ischemia may contribute to adverse transplant outcomes. Of note, we identified upregulation of the oxygen dependent oxidative phosphorylation (OXPHOS) pathway with increasing duration of cold ischemia injury within the oxygen-lean inner medulla region of the kidney. The inner medulla prefers the use of glycolytic pathways to meet its energy demands and is usually not accessible by biopsy (*18*, *19*). As such, we believe the cold ischemia-associated molecular presentation within the inner medulla is atypical. We also compared the transcriptional trends between the corresponding kidney compartments subject to cold ischemia injury with that of published data on warm ischemia injury (ischemia-reperfusion injury) which is relatively well characterized (*20*). We observed *Spp1* to be upregulated within the all the three compartments within both cold and warm ischemia tissue. In contrast, the OXPHOS pathway was observed to be upregulated within the inner and outer medulla of kidney tissue subjected to cold ischemia injury while it was downregulated during warm ischemia tissue, highlighting both covarying and divergent molecular processes between these two injury models. We believe these biological insights may help towards understanding why prolonged cold ischemia injury is associated with worse clinical outcomes. Moreover, these observations point towards a gradual pathophysiologic and adaptive changes within this deeper kidney tissue region with increasing cold ischemia time, highlighting the need for increased attention to the inner medulla of kidney allografts. An interactive browser with our data is available to facilitate exploration at https://jef.works/CellCarto-ColdIschemia/.

**Figure 1.**
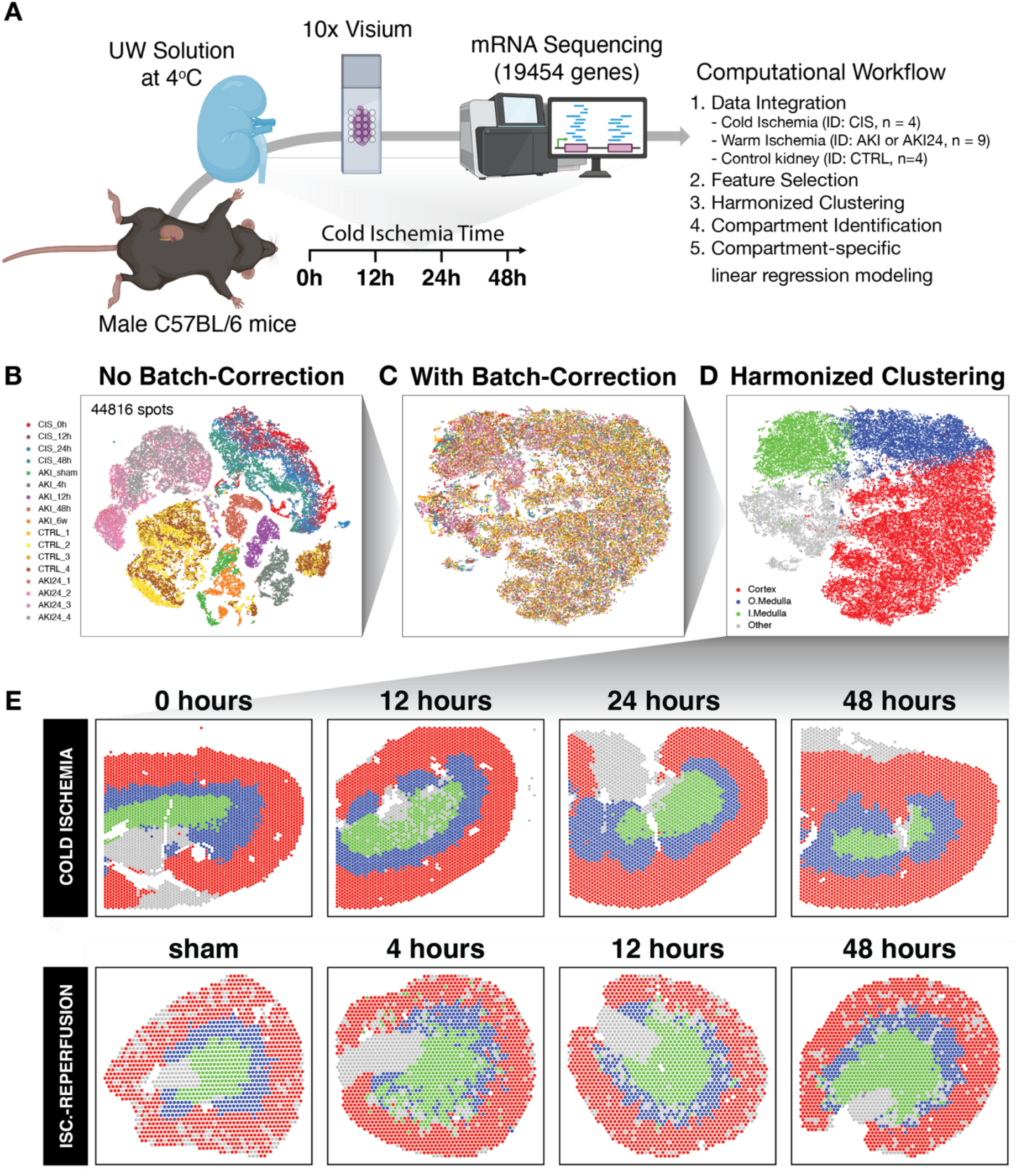
Computational workflow for analysis of multiple murine kidney spatial transcriptomics datasets. **(A)** Spatial transcriptomics datasets were generated from murine kidneys subjected to cold ischemia injury of various durations: 0, 12, 24 and 48 hours. Thereafter, the datasets were integrated with previously published ST datasets for downstream analysis. **(B)** tSNE plot demonstrating sample-specific effects within the integrated ST datasets. **(C)** Harmonized tSNE plot leading to a shared embedding across all datasets. **(D)** Clustering facilitated identification of shared transcriptionally distinct compartments across all the datasets, namely cortex, outer medulla (O.Medulla), inner medulla (I.Medulla) and other. **(E)** Spatial plots visualizing the anatomic location of the different transcriptionally-defined compartments within the kidney tissue specimens belonging to different experimental groups like cold ischemia injury and ischemia-reperfusion injury. Created with BioRender.com

## Results

### 1. Computational workflow identified shared spatial compartment across multiple murine kidney spatial transcriptomics datasets

To establish a quantitative molecular understanding of cold ischemia across the entire kidney, we applied 10x Visium to perform spot-resolution full transcriptome spatial transcriptomic profiling of cold ischemia kidney injury in a murine model. We applied 0, 12, 24 and 48 hours of cold ischemia injury that represent different durations of cold ischemia in a deceased-donor kidney transplant setting (Fig. 1A, S1A-D) (*21*). Quality control filtering resulted in 19,454 unique gene species detected across 12,530 spots across the cold ischemia kidney injury tissue specimens (4.18 ± 0.29 transcripts/spot and 2.6 ± 0.96 spots/gene (log_10_ scale)) (Fig. S1E-L). This raw gene count matrix was further CPM (counts per million) normalized to control for potential technical variations in sequencing depth across spots. To facilitate comparative analysis across the cold ischemia kidney specimens as well as with other murine kidney specimens from previously published studies (*20*, *22*), including healthy control kidneys as well as kidneys with warm ischemia-reperfusion injury (also a model native kidney acute kidney injury-AKI), we sought to computationally integrate all 17 ST datasets (44,816 spots) to identify transcriptionally similar tissue compartments via graph-based clustering (Fig. 1B). However, such unified clustering was driven by batch effects as well as unique biological processes resulting in condition and sample-specific clusters, thereby hindering the identification of shared features. We therefore performed batch correction using Harmony (17) (Fig. 1C) and re-clustered to identify shared compartments across all datasets (Fig. 1D). Due to lack of single cell resolution and concerns for variations in cell-type proportions confounding small clusters, we opted for coarser clustering. Ultimately, this computational workflow led to identification of distinct compartments which we labeled as cortex, outer medulla (O.Medulla), inner medulla (I.Medulla) based on their relative anatomical location within all the kidney tissue specimens (Fig. 1E, Fig. S1M,N). Review of the corresponding H&E images by an expert kidney pathologist validated our labels, confirming congruence between the transcriptional and histological assessments. Remaining tissue which was not part of the previous three compartments was labeled as “other,” likely corresponding to the transition from medulla to pelvis based on the pathologist’s evaluation and was not used for any analyses henceforth. We were then able to perform differential expression analysis across compartments of all ST datasets (Fig. S2) to identify marker genes of the different compartments, which were found to be consistent with known annotations of associated functional units. For example, markers for functional units like *Slc14a2* for descending thin limb of Loop of Henle (LOH) and *Slc5a3* for thick descending limb of LOH (*24*) were identified as one of the top gene markers for inner medulla compartment. Similarly, proximal tubule S3 segment marker *Slc22a7* (*25*) was identified as one of the top gene markers for outer medulla while pan-proximal tubule marker *Slc34a1* and proximal convoluted tubule marker *Slc5a2* were associated with the cortex (18) (Fig. S3). This observation was consistent across all datasets. As such, our computational workflow identified spatially distinct compartments, thereby enabling us to further analyze transcriptomic temporal dynamics in the different compartments of kidney tissue.

### 2. Compartment-specific analysis characterized spatially confined gene expression dynamics in cold ischemia kidney injury

To characterize the temporal transcriptional dynamics associated with cold ischemia injury, we leveraged linear regression modeling to identify genes with normalized expression that significantly change with duration of cold ischemia injury (i.e., CIT) within each identified kidney compartment. First, across all compartments, we observed that the total mRNA counts per spot decreased over the duration of cold ischemia injury (0-48hours), which might be due to mRNA degradation in an ischemic tissue environment but might also be attributable to technical variations in sequencing depth (Fig. S4A-C). To control for such potential technical variation, we performed counts-per-million (CPM) normalization thereby adjusting raw counts to a proportional quantification with a common sequencing depth. We then applied linear regression to each normalized gene for spots in each compartment separately to identify significantly trends based on multiple-testing corrected p-values and regression R^2^ filters (Methods). Such linear regression modeling identified many genes that appear downregulated (proportionally decreased) with longer CIT i.e., exhibited a significant downward temporal trend (number of genes in inner medulla: 494 (2.5%), outer medulla: 562 (2.9%) and cortex: 492 (2.5%))(Table S1).Other genes appear upregulated (proportionally enriched) with longer cold ischemia time i.e., exhibited a significant upward temporal trend (inner medulla: 305 (1.6%), outer medulla: 471 (2.4%) and cortex: 531 (2.7%))(TableS2). Comparing across compartments, we found temporally upregulated genes either in a compartment-agnostic or compartment-specific manner. For compartment-agnostic trends, 128 (0.66%) genes exhibited an upward temporal trend across all compartments whereas 136 (0.7%) genes exhibited a downward temporal trend across all compartments (Table S3). For example, *Ig8p7* (Fig. 2A), *Fth1,* and *Spp1* were upregulated over time across all compartments whereas *Kap* (Fig. S5A)*, Keg1,* and *Acsm2* were downregulated over time across all compartments. We also identified genes that exhibited a compartment-specific temporal trend (upward trend: inner medulla: 102 (0.52%), outer medulla: 164 (0.87%), cortex: 211 (1.1%)(Table S4); downward trend: inner medulla: 249 (1.28%), outer medulla: 169 (0.86%), cortex: 150 (0.77%) (Table S5)). For example, *Ranbp3l* was uniquely upregulated over time in the inner medulla (Fig. 2B), whereas *Ttc36* was uniquely upregulated in the outer medulla (Fig. 2C), and *Slc34a1* in the cortex (Fig. 2D). In contrast, *Egf* was uniquely downregulated over time in the inner medulla, whereas *Gpx1* was uniquely downregulated in the outer medulla, and *Chpt1* in the cortex (Fig. S5B-D). Within each compartment, we observed a great variation in the magnitude of gene expression changes as quantified by linear regression slopes, with most genes demonstrating subtle changes as reflected in small slopes (normalized scores<0.5) while a few genes exhibited marked larger, outlier slopes such as *Fth1, Ig8p7*, and *Kap*. (Fig. 2E, F-H, Table S1,2). To investigate whether these trends covary between compartments, we compared the regression slopes of genes in the different compartments. Pearson correlation analysis suggested that the cortex and outer medulla (Pearson correlation coefficient, PCC= 0.98, p<0.05, R^2^= 0.96) (Fig. 2F, Table S6,7) are more similar to each other in terms of temporal gene expression changes as compared to the inner medulla (cortex vs inner medulla: PCC= 0.66, p<0.05, R^2^= 0.43) (Fig. 2G, Table S6,7); outer medulla vs inner medulla: PCC= 0.71, p<0.05, R^2^= 0.51) (Fig. 2H, Table S6,7)). As such, our compartment-specific temporal analysis identified compartment-agnostic as well as compartment-specific gene expression trends.

**Figure 2.**
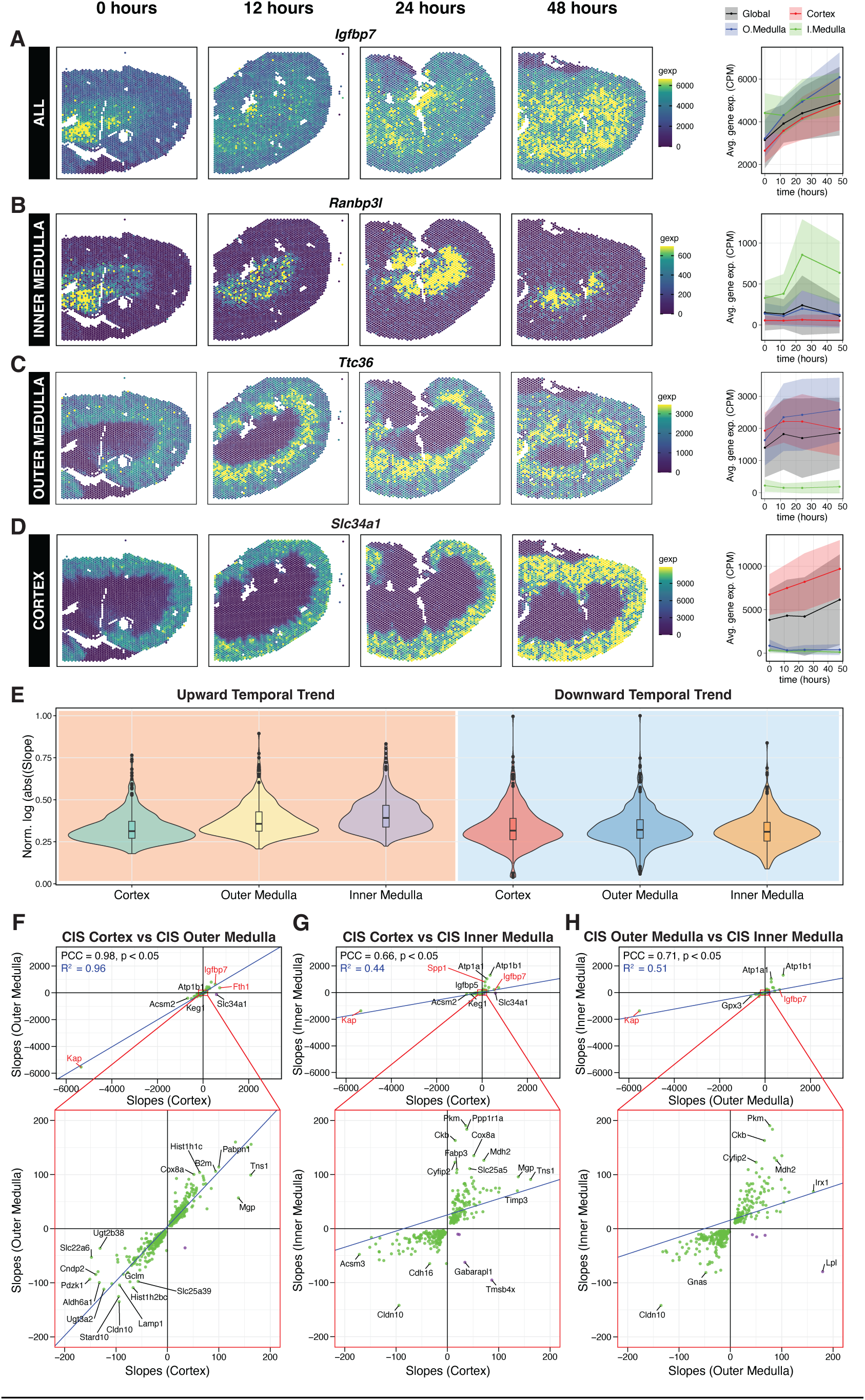
Compartment-specific temporal transcriptional dynamics in cold ischemic kidneys. **(A)** Spatial gene expression plots and corresponding ribbon plots (mean ± s.d) demonstrating spatiotemporal dynamics of representative genes that demonstrated a significant upward temporal trend during the cold ischemic phase (0-48 hours). For example, **(A)** *Ig8p7* was significantly temporally upregulated in all the compartments with varying degrees whereas **(B)** *Ranbp3l,* **(C)** *Ttc36* and **(D)** *Slc34a1* were uniquely, significantly temporally upregulated in the inner medulla, outer medulla and cortex compartments of cold ischemic injury kidney tissue respectively. **(E)** Violin plots highlight the distribution of magnitude of transcriptional changes (linear regression slopes) within the different compartments of CIS kidney tissue. Scatterplots depicting the correlation analysis of the temporal transcriptional dynamics between the different compartments: **(F)** cortex vs outer medulla, **(G)** cortex vs inner medulla and **(H)** outer medulla vs inner medulla of the CIS kidney tissue. Genes exhibiting covarying (green) or divergent (purple) temporal trend within these compartments can be visualized along with their Pearson correlation coefficients (PCC) and linear regression slopes (blue line). Select genes exhibiting strong temporal changes within these compartments have been highlighted red.

To gain insight into whether such spatiotemporal gene expression changes may be attributable to changes in cell-type composition of the kidney tissue during cold ischemia injury, we performed reference-free cell type deconvolution to identify 7 major cell types and quantify their proportions across time points (Fig. S4E). Unlike transcriptional changes, cell type composition remained generally consistent even with longer cold ischemia time, especially within the inner medulla. (Fig. S4F). Subtle compartment specific changes were observed for deconvolved cell type 2, whose population reduced within the cortex and outer medulla. Based on the associated topmost upregulated marker genes and enriched (GO) pathways (Table S8-9), we interpret this deconvolved cell-type 2 to be proximal tubule cell, which is known to decrease in proportion under other ischemic conditions such as acute kidney injury (*26*). Nevertheless, these subtle changes in cell-type proportions cannot fully explain the strong compartment-specific transcriptional changes, particularly in the inner medulla.

### 3. Pathway characterization of identified spatiotemporal trends highlight elevated oxidative phosphorylation within the inner medulla

To better understand the molecular pathways associated with genes that demonstrated significant time dependent expression with increasing CIT, we performed gene set enrichment analysis using KEGG and HALLMARK (HM) annotations on the significantly temporally upregulated and temporally downregulated genes with longer CIT obtained through our previous linear regression analysis ranked by slope in a spatial compartment specific manner (Methods). We identified several significantly enriched molecular pathways and evaluated whether they exhibited covarying or divergent trends across different compartments based on their normalized enrichment scores (NES) (Fig. 3A-C, Table S10, S11). Within the kidney inner medulla, we observed significant positive enrichment (NES>0) of energy metabolism and neurodegenerative pathways amongst several enriched biological pathways (Fig. 3A), suggesting that genes within these pathways were being temporally upregulated in this compartment with increasing CIT. For example, the metabolic pathway for oxidative phosphorylation (OXPHOS) exhibited a strong (NES=4.47 (KEGG), 5.06 (HM)) to moderate (NES=2.29 (KEGG), 2.13 (HM)) positive enrichment within the inner and outer medulla respectively, while exhibiting a negative enrichment (NES=-2.56 (HM)) within the cortex (Fig. 3A-C, D-F, Table S10). In conjunction, upstream metabolic pathways that help facilitate OXPHOS like citrate cycle (TCA cycle) (NES=-2.07) and fatty acid degradation (NES=-2.67) also exhibited negative enrichment within the cortex (Fig. 3C, Table S10). Within the inner medulla, the leading-edge genes of the OXPHOS pathway were related to various processes of mitochondrial machinery like *Uqcr11* (complex III) and *Cox6a1* (complex IV), (Fig. 3 G,H). Therefore, to validate these spatiotemporal pathway trends in a larger cohort, we performed compartment-specific qPCR using 4-5 animals per time point for two time points (CIS 0hours vs CIS 48 hours) (Methods). Our qPCR analysis demonstrated similar upward transcriptional trends for *Uqcr11* (Fig. 3I) and *Cox6a1* (Fig. 3J) within the kidney medulla as observed in spatial transcriptomics.

**Figure 3.**
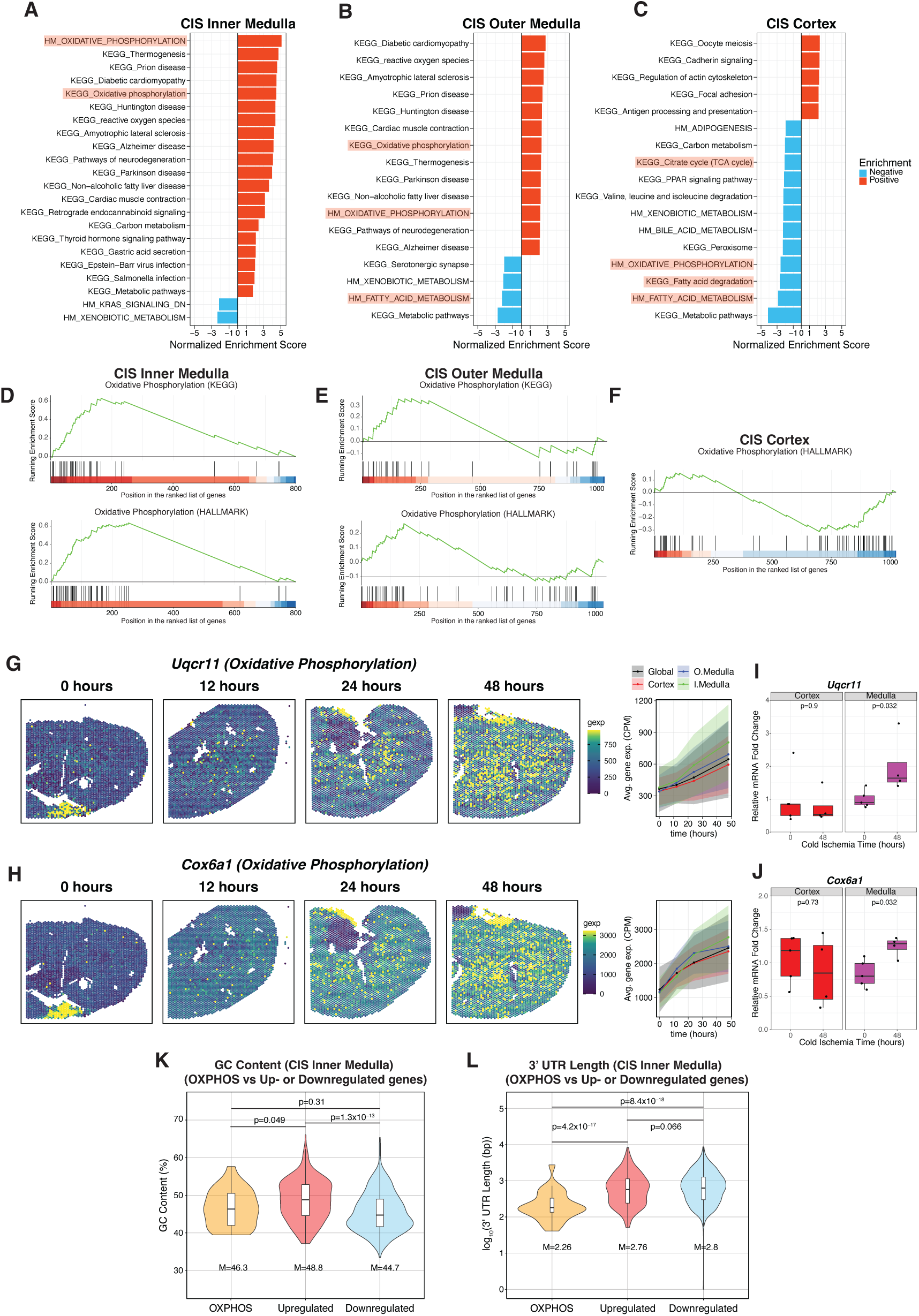
Compartment-specific enrichment of molecular pathways within cold ischemic kidney tissue. Bar plots highlighting all the enriched molecular pathways (KEGG or HALLMARK (HM)) that demonstrated similar temporal dynamics (enrichment: positive (red), negative (cyan)) within the corresponding **(A)** inner medulla **(B)** outer medulla and **(C)** cortex of cold ischemic injury kidney tissue. (Prominent energy metabolism pathways like oxidative phosphorylation and upstream metabolic pathways that facilitate OXPHOS like fatty acid metabolism and TCA cycle have been highlighted within the list of all enriched pathways.) Running enrichment score plots highlighting differential enrichment of molecular pathways related to oxidative phosphorylation within the **(D)** inner medulla, **(E)** outer medulla and **(F)** cortex of the CIS kidney tissue. Spatial gene expression plots and corresponding line plots demonstrating spatiotemporal dynamics of representative genes associated with the above enriched energy metabolism related pathways: **(G)** *Uqcr11* and **(H)** *Cox6a1* (oxidative phosphorylation). Quantitative comparison of qPCR derived mRNA transcript levels of **(I)** *Uqcr11,* **(J)** *Cox6a1* between the cortex and medulla (inner medulla + outer medulla) of CIS kidney tissue. (n=5 for 0 hours (cortex and medulla samples each), n=4 for 48 hours (cortex and medulla each). **(K)** Violin plots comparing distribution of GC content of OXPHOS related genes and all the temporally upregulated and downregulated genes within the CIS inner medulla. **(L)** Violin plots comparing distribution of log_10_ transformed 3’ UTR length (bp: base pairs) of OXPHOS related genes and the temporally upregulated and downregulated genes within the CIS inner medulla. M denotes the median values for the corresponding violin plots in figures **K** and **L**. Two-sided Wilcoxon rank sum test was used for statistical testing in figures **K** and **L**. Benjamini-Hochberg method was used for multiple hypothesis testing for gene set enrichment analysis.

This observation of temporal upregulation of genes related to OXPHOS within the inner medulla was unexpected because the inner medulla in a normal kidney, being in an oxygen-lean environment, is known to prefer the anaerobic glycolytic pathway (*27*). Given that we previously observed the total mRNA counts to decrease over the duration of cold ischemia injury (Fig. S4A-C), we sought to assess whether this identified trend could be due to normalization. Therefore, we further evaluated the raw counts of OXPHOS genes (leading edge genes from gene set enrichment analysis from HALLMARK pathway, Fig. 3D-F). Raw counts demonstrated a similar declining trend within the outer medulla (slope=-4.7) and cortex (slope=−9.9) but was relatively stable within the inner medulla (slope=−0.38) (Fig S4D-F), thereby confirming the compartmentally-distinct temporal trends identified through our previous analysis. As such, the ratio of OXPHOS mRNA count to total mRNA count increased strongly in the inner medulla (slope=0.042) and moderately in the outer medulla (0.017) while it declined in the cortex (slope=−0.011) (Fig. S4G-I). This suggests that the temporal upregulation (positive regression slope) of OXPHOS genes within the inner medulla and outer medulla and downregulation (negative regression slope) within the cortex with increasing cold ischemia injury identified by our linear regression analysis on the CPM normalized gene expression might be better interpreted as a proportional enrichment and a proportional depletion respectively.

Next, we interrogated whether the proportional enrichment of OXPHOS genes could stem from relative stability of mRNA molecules. Genes with lower GC content and shorter 3’ UTR length (base pairs) are known to be relatively more stable (*28*). Consequently, they could degrade slower in an ischemic environment relative to other genes, resulting in a proportional enrichment and perceived temporal upregulation over time due to degradation of the other genes alone. If temporal upregulation was due to higher relative mRNA stability, we would anticipate temporally upregulated genes to have the lower GC content and shorter 3’UTR length compared to temporally downregulated gene transcripts. To test this, we compared the GC content and 3’UTR lengths of the OXPHOS genes with the other temporally upregulated and downregulated genes within the inner medulla. We found that the GC content of the OXPHOS genes (median=46.3%) was very comparable to that of all the other temporally upregulated (median=48.8%) and downregulated (median=44.7%) genes within the inner medulla (Fig. 3K). Further, while other temporally upregulated and downregulated genes exhibited a significant difference in GC content, the direction of the difference was contrary (higher in upregulated genes) to what we would expect if temporal upregulation was due to higher relative mRNA stability. In contrast, the 3’UTR length (i.e., log_10_ transformed 3’ UTR length) of these upregulated (median=2.76) and downregulated genes (median=2.8 base pairs) within the inner medulla were comparable (Fig. 3L). Taken together, these findings suggest that the temporal upregulation of genes within the inner medulla is likely not solely attributable to higher relative mRNA stability, and that other mechanisms could potentially contribute to this observed transcriptional trend with increasing duration of cold ischemia injury. Still, the 3’ UTR lengths of OXPHOS genes (median=2.26) was significantly lower than that of all other temporally upregulated and downregulated genes, consistent with enhanced mRNA stability (Fig. 3L). However, again, the lack of consistency based on GC content and lack of difference in 3’ UTR length for other upregulated and downregulated genes suggest that this characteristic alone may be insufficient to fully explain the observed temporal patterns.

### 4. Compartment-specific overlapping and divergent molecular dynamics identified between cold ischemia and warm ischemia-reperfusion-induced AKI

To better understand the pathogenesis of kidney cold ischemic injury, we sought to examine its spatiotemporal transcriptomic dynamics alongside another ischemic injury model to identify potential shared molecular processes. We therefore repeated the spatiotemporal analysis with a set of previously published ST datasets from a well-characterized murine model of warm ischemia-reperfusion-induced acute kidney injury (AKI) at various timepoints postinjury covering the peak phases of injury response (sham to 48 hours)(*20*). Our temporal analysis recapitulated putative AKI injury markers like *Havcr1, Lcn2 and Spp1* with *Spp1* demonstrating the sharpest change (linear regression slope) during the acute injury phase (Fig. S6A-C), further highlighting the ability of our spatiotemporal analysis to detect known injury-related transcriptional changes. Cross-compartmental analysis highlighted similar temporal transcriptional dynamics between the AKI cortex and inner medulla (PCC=0.87, p<0.05, R^2^=0.75) while the outer medulla and inner medulla were comparatively less similar (PCC=0.46, p<0.05, R^2^=0.21) (Fig. S6D-F).

We next compared the temporal transcriptional dynamics between the corresponding compartments of the two injury models (i.e., CIS vs AKI). Compared to CIS, in the warm ischemia-reperfusion-induced AKI model, while compartment-specific correlation analysis indicated a weak correlation between the inner medulla (PCC=0.09, p>0.05; R^2^∼0 (0.007)) (Fig. 4A), comparatively stronger correlations were observed between the outer medulla (PCC=0.84, p<0.05; R^2^=0.7) (Fig. 4B) and cortex (PCC=0.53, p<0.05; R^2^=0.26) (Fig. 4C) of both the injuries. Interestingly, we observed *Spp1* upregulation across all compartments in both injury models with the inner medulla exhibiting the largest expression (Fig. 4A-C). Since *Spp1* expression is strongly temporally upregulated within all compartments of AKI kidney tissue, the similar trend of temporal upregulation of *Spp1* within all compartments of CIS kidney tissue might be suggestive of a pathological environment within the CIS tissue.

**Figure 4.**
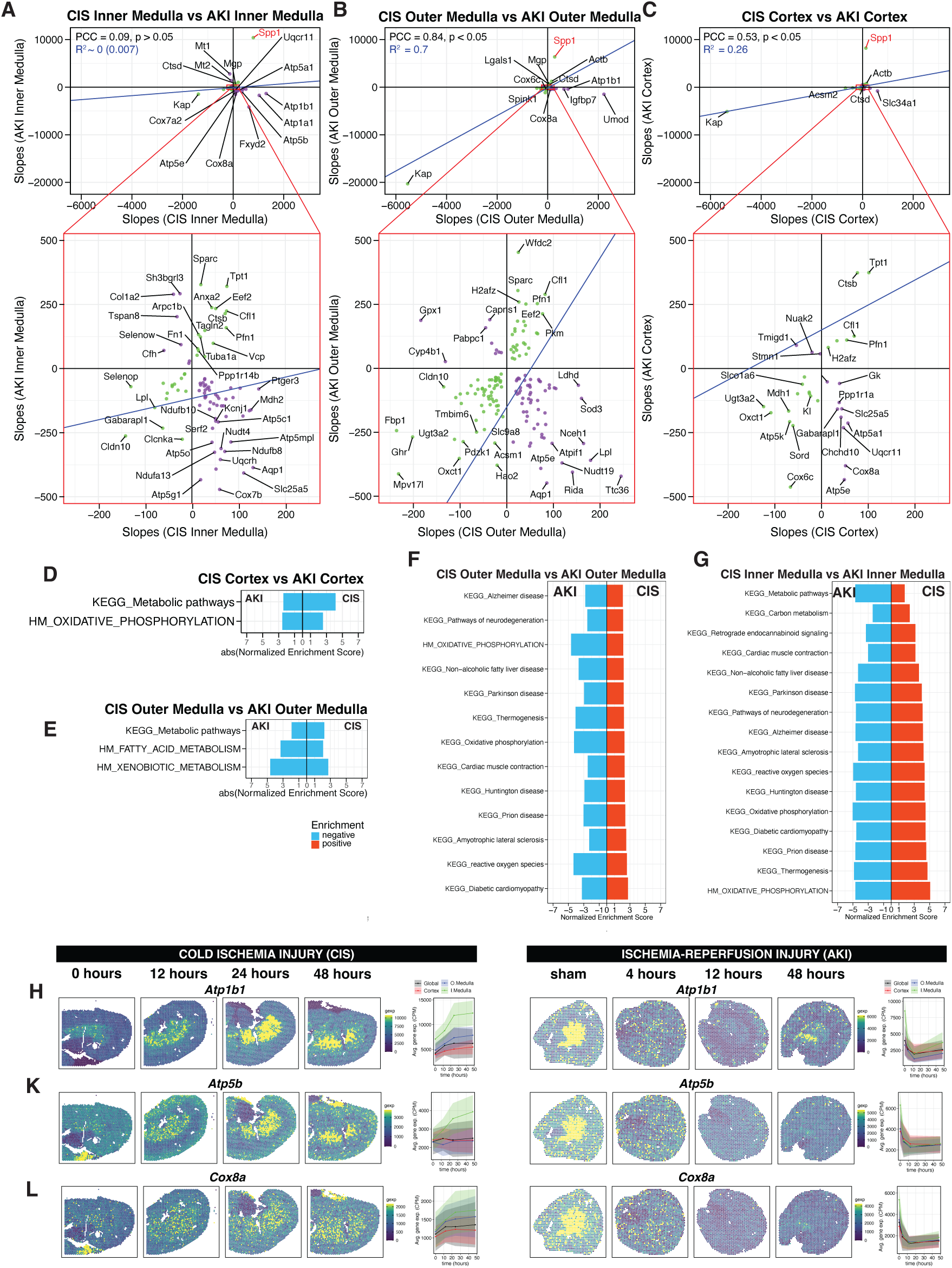
Compartment-specific comparison of transcriptional dynamics between the CIS and AKI kidney tissue. Scatter plots depicting the correlation analysis of the temporal transcriptional dynamics between the corresponding **(A)** inner medulla, **(B)** outer medulla and **(C)** cortex tissue compartments of the two kidney injury models i.e., cold ischemic injury (CIS) and warm ischemia-reperfusion injury (AKI) kidney tissue. Genes exhibiting covarying (green) or divergent (purple) temporal trend within these compartments can be visualized along with their Pearson correlation coefficients (PCC) and linear regression slopes (blue line). Bar plots highlighting all the enriched molecular pathways that demonstrated similar temporal dynamics (enrichment: positive (red), negative (cyan)) within the corresponding **(D)** cortex, and **(E)** outer medulla, and divergent temporal dynamics within the corresponding outer medulla and inner medulla of CIS and AKI kidney tissue. Spatial gene expression plots and corresponding ribbon plots demonstrating spatiotemporal dynamics of genes associated with OXPHOS pathway like **(K)** *Atp1b1,* **(L)** *Atp5b,* and **(M)** *Cox8a* within both the CIS and AKI kidney tissue. Benjamini-Hochberg method was used for multiple hypothesis testing for gene set enrichment analysis.

Based on these observations, we performed gene set enrichment analysis to characterize pathways impacted in AKI in a compartment-specific manner as we did with CIS (Fig. S7, Table S12, S13). For the covarying pathways across compartments within the AKI tissue, pathways associated with energy metabolism like oxidative phosphorylation, citrate cycle (TCA cycle), fatty acid degradation, etc., demonstrated negative enrichment across compartments. Positive enrichment of pathophysiologic processes like epithelial-to-mesenchymal transition (or EMT) (ID: EPITHELIAL_MESENCHYMAL_TRANSITION; genes: *Spp1, Col1a1, Lgals1, Vim*, etc.) was observed within the all the three compartments of AKI kidney tissue, suggesting the potential development of a process leading to wound healing and fibrosis. Of note, positive enrichment of coagulation pathway (ID: COAGULATION; genes: *Clu, Timp1, Mmp2,* etc.) within the inner medulla and complement pathway (ID: COMPLEMENT; genes: *CR, Pfn1, Ctsd, Timp1*, etc.) within the inner and outer medulla hinted at potential endothelial dysfunction/vascular remodeling and immune-complex formation activating the complement (classical) pathway during reperfusion. Processes like EMT, complement activation and microvascular dysfunction known to be associated with ischemia-reperfusion injury (*29*) validate the ability of our computational workflow in identifying relevant temporal gene and pathway changes that accompany the pathogenesis of diseases like AKI.

We also compared the enriched pathways between the two injury models in a compartment-specific manner (Fig. 4 D-G, Table S14, S15). Compartment-specific pathways exhibiting divergent trends between the two injury models were mostly related to energy metabolism and neurodegenerative-linked molecules and were found to be confined to the inner and outer medulla. For example, the OXPHOS pathway was upregulated only within the inner medulla and outer medulla of CIS while it was downregulated within that of AKI (Fig. 4F, G). This finding is consistent with the weak correlation observed between the CIS vs AKI inner medulla changes mentioned earlier (Fig. 4A).

Acknowledging that the AKI and CIS datasets were derived from female and male C57BL/6 mice respectively, we next sought to evaluate whether sexual dimorphism could be contributing to these compartment-specific divergent transcriptional dynamics. We therefore analyzed normal control kidneys (CTRL) and 24-hours warm ischemia reperfusion injury kidney tissues (AKI24) from male C57BL/6 mice (*22*). First, we observed that the presentation of AKI marker genes like *Havcr1*, *Lcn2* and *Spp1* in the 24-hour male (AKI24) kidney tissues was very similar to that of female kidney specimens (AKI) at 12 and 48 hours (Fig. S6 A-C, G-I). Next, we performed differential expression analysis between inner medulla compartment belonging to the CTRL (male) and AKI24 (male) kidneys. We observed that the leading genes: associated with OXPHOS, the divergent pathways identified earlier (CIS inner medulla vs AKI inner medulla), exhibited similar expression trend during AKI injury within the inner medulla of these male kidneys as what was previously identified in the female kidneys. In particular, OXPHOS related genes like *Atp1b1, Atp5b and Cox8a* previously identified to be downregulated over time within the inner medulla in AKI in the female kidneys were again identified as downregulated within the inner medulla of AKI24 compared to CTLR in the male kidneys (Fig. S8A). We performed gene set enrichment analysis on the differentially expressed genes between the corresponding compartments of CTRL and AKI (24 hours) kidneys in male and observed negative enrichment of OXPHOS pathway within inner and outer medulla while no enrichment in cortex was observed (Fig. S8B). The above observations highlight that, overall, the spatiotemporal transcriptional dynamics for the OXPHOS pathway observed within male and female AKI kidney specimens are consistent and thus the divergent temporal trends in OXPHOS gene expression between CIS and AKI kidney specimens are likely not driven by solely sexual dimorphism. Collectively, the transcriptomic presentation of both CIS and warm IRI (AKI) display some molecular similarities like the upregulation of *Spp1* within all the compartments highlighting potential similarities in injury response across these distinct ischemic models. While divergent temporal trends like OXPHOS within the inner medulla highlight that while both injuries share commonalities, their compartment-specific metabolic response to injury may differ.

### 5. Compartment-specific atypical metabolic changes may be implicated with cold ischemia injury

Our spatiotemporal transcriptomic analyses highlighted a temporal upregulation of the oxygen demanding OXPHOS pathway with increasing duration of cold ischemia injury within the oxygen-lean inner medulla of the kidney tissue. The medullary region of the kidney is known to prefer glycolysis for meeting its energy requirements (*18*, *19*). Thus, we checked for glycolysis related genes within the kidney tissue. Interestingly, we observed temporal upregulation of glycolysis related gene *Pkm* within all compartments, with moderate upregulation within the outer medulla (regression slope=76.77) and cortex (regression slope=35.99) while the strongest upregulation (regression slope=190.48) was observed within the inner medulla (Table S2, Fig 5A). Similarly, we observed the spatiotemporal expression of glucose transporter gene *Slc2a1* (GLUT1) mirror that of *Pkm*, with the strongest upregulation within the inner medulla (regression slope=70.22) and moderate upregulation within the outer medulla (regression slope=20.32) and cortex (regression slope=10.25) (Table S2, Fig 5B).

**Figure 5.**
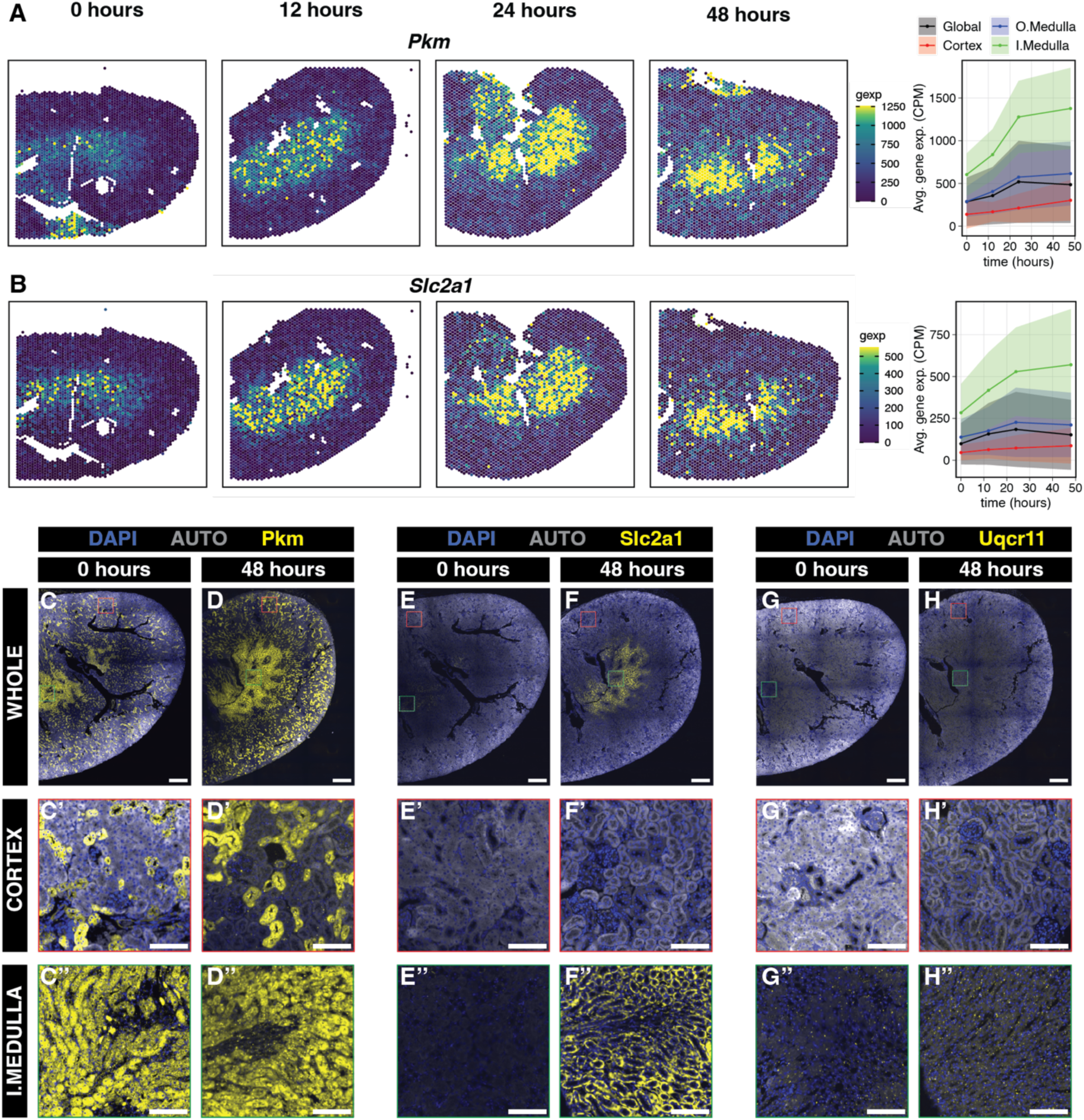
Experimental validation of spatiotemporal transcriptomic trends using immunofluorescene. Spatial gene expression plots and corresponding ribbon plots demonstrating spatiotemporal dynamics of **(A)** *Pkm* and **(B)** *Slc2a1* within cold ischemia kidney injury tissue. Immunofluorescence images of kidney tissue cross-sections displaying protein level spatiotemporal expression of **(C,D)** Pkm, **(E,F)** Slc2a1 (Glut1) and **(G,H)** Uqcr11 at 0- and 48 hours of cold ischemia injury respectively. Two representative regions (400*μ*m × 400*μ*m each) highlighted across all tissue sections belonging to the cortex (red) and inner medulla (green) region. **(C’-H’, C’’-H’’)** Magnified images of the representative cortex and inner medulla regions for Pkm, Slc2a1 and Uqcr11 respectively. Pseudo-colors used: blue (DAPI), gray (autofluorescence (AUTO)) and yellow (protein marker of interest). Scale bar: 500 *μ*m **(C-H)**, 100 *μ*m **(C’-H’,C’-H’’)**.

To further validate these spatiotemporal transcriptional trends, we performed immunostaining to visualize the protein expression patterns for Pkm, Slc2a1, and Uqcr11 across kidney tissue regions for 0- and 48- hours cold ischemia time points (Fig. 5C-H’’). Qualitative evaluation of the immunostained cold ischemic tissue specimens demonstrated the expected subcellular and tissue-level spatial molecular patterns, with cytosolic expression of Pkm, membrane expression of Slc2a1, and mitochondrial expression of Uqcr11 being largely localized to the inner medulla (Fig. 5). Further, we observe sustained protein level expression of Pkm (Fig. 5C-C”,D-D”) and a marked increase in protein level expression of Slc2a1 (Fig. 5E-E”,F-F”) and Uqcr11 (Fig. 5G-G”,H-H”) particularly within the inner medulla between 0 and 48 hours cold ischemic kidney tissues. These results indicate a general spatiotemporal molecular agreement at both at the transcriptomic and proteomic level with higher expression of Pkm, Slc2a1 and Uqcr11 within the inner medulla as compared to the cortex, particularly after 48h of cold ischemia. These compartment-specific temporal trends in both gene and protein level changes suggest an overall atypical metabolic presentation that accompany cold ischemia injury in kidneys (Fig. 6).

**Figure 6.**
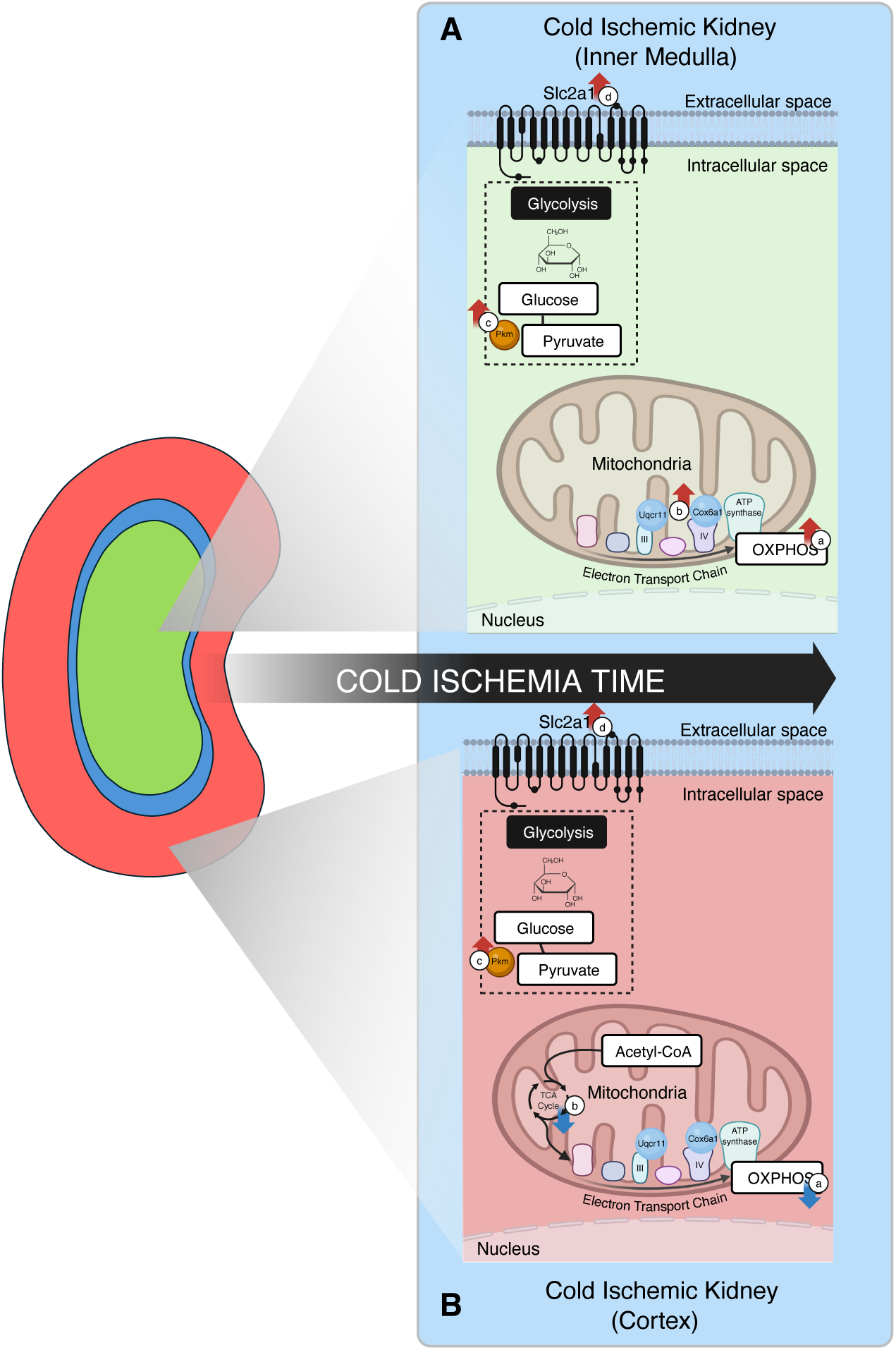
Cold ischemia injury induced atypical compartment-specific metabolic changes in murine kidneys. Atypical compartment-specific metabolic changes that accompany cold ischemia injury in kidneys: **(A)** Cold Ischemic Kidney (Inner Medulla): During cold ischemia injury, molecular pathways related to energy metabolism like (a) OXPHOS and was positively enriched (red arrows) within the inner medulla. (b) OXPHOS related genes *Uqcr11* and Cox6a1 related to complex III and IV respectively of mitochondrial electron transport chain were upregulated. Strong upregulation of (c) glycolysis related gene of *Pkm* (transcriptomic) expression and (d) glucose transporter gene *Slc2a1* (both transcriptomic and protein level) within the inner medulla. **(B)** Cold ischemic kidney (cortex): Metabolic pathways like (a) OXPHOS and (b) TCA cycle were negatively enriched (blue arrows) during cold ischemia injury. Finally, (c) glycolysis related gene *Pkm* and (d) glucose transporter gene *Slc2a1* were observed to be weakly upregulated within the cortex (transcriptomic level only). (Different colors represent different compartments of the kidney tissue: inner medulla (green), outer medulla (blue) and cortex (red)). Created with BioRender.com.

## Discussion

Long durations of static cold preservation storage results in cold ischemia injury and contributes to worse graft outcomes (*6*). However, the molecular events during the pathogenesis of this injury are incompletely understood, particularly in the medulla of the kidney, which is less amenable to safe percutaneous or wedge biopsy sampling in a clinical setting. We therefore performed a full transcriptome spatial characterization of cold ischemia in murine kidneys (0-48 hours) using ST with 10x Visium. Using a data driven computational analysis approach, we identified temporally varying genes within transcriptionally distinct compartments of the kidney tissue, enabling compartment-specific investigation of gene and pathway dynamics.

Through time-resolved analysis of spatial molecular profiles across kidney compartments, our analysis identified genes and pathways that demonstrated compartment-specific, time-dependent proportional enrichment and depletion as significant temporal upregulation and downregulation respectively with increasing duration of cold ischemia injury. These genes were associated primarily with energy metabolism molecular pathways identified through gene set enrichment analysis. Particularly, temporal upregulation of the OXPHOS pathway within the oxygen-lean kidney inner medulla, which typically relies on the anaerobic glycolytic pathway. As such the molecular presentation within the inner medulla is atypical. This proportional enrichment is likely not explained by cell type proportion changes or relative mRNA stability alone. Comparative analysis of cold and warm ischemia kidney tissue identified that OXPHOS was uniquely upregulated within the inner medulla of cold ischemia kidney tissue while it was observed to be temporally downregulated within the inner medulla of warm ischemia-reperfusion injury kidney tissue. Overall, the identified trends with increasing cold ischemia time suggest heightened OXPHOS within the inner medulla and further investigation might be necessary to evaluate whether this is a pathological development or a compensatory mechanism.

Although our current study brings in new molecular insights into the pathogenesis of cold ischemic injury within the kidney, there are limitations which may be addressed in future studies. First, the lack of single cell resolution in our spatial transcriptomic profiling limits our understanding of the cell-type specific effects within the different compartments during cold ischemia injury. For example, although we observe changes in OXPHOS gene expression within the inner medulla, whether this change is driven by a specific cell-type like collecting duct cells or multiple cell-types within that region remains unclear. Additionally, resolution limits our ability to perform characterization of specific functional tissue units like the glomerulus which has clinical relevance in the context of an ischemic setting. Single-cell resolution ST platforms like Xenium (*30*) or MERFISH (*31*) could be utilized to perform such cellular or glomerulus-specific characterization, though they are currently limited in the number of genes they can profile simultaneously and therefore can be informed by full-transcriptome characterizations such as this study. Alternatively, compartment-specific single-cell RNA-sequencing could also be used, though its spatial fidelity is limited by the precision of microdissection, making compartment-specific analyses challenging, particularly in the outer medulla, which is anatomically interposed between the cortex and inner medulla. Nevertheless, we anticipate that the compartment-specific insights from our ST analysis will inform future compartment-specific single-cell investigations. In addition, although we analyzed ∼3133 spatially distinct spots per sample, spots were derived from same tissue section. Considering the limited number (i.e., n=1) of biological replicates per time point, we decided to focus our analysis only on consistent transcriptomic trends during the entire duration of cold ischemia injury (0-48 hours). We did not analyze any time-point specific transcriptomic changes. Further studies with increased biological replicates would enable us to characterize early vs late response and enhance our understanding of the injury progression. Although we acknowledge that lack of biological replicates limits the generalizability of exact regression slopes, we anticipate the general ranks and signs of temporal trends across genes to be more robust, which would in turn lead to robust gene set enrichment results. As the high cost of ST technologies to generate increased sample sizes remains a limiting factor, we employed qPCR with a larger animal cohort (n=4-5 per timepoint) to check the validity of key findings from ST as well as immunostaining to establish concordance at the protein level. Application of our computational workflow to AKI ST datasets further recapitulated known molecular trends related to warm ischemia kidney injury and consistent between male and female animals, underscoring the general validity of our analytical pipeline and the robustness of identified general trends despite being limited in the number of biological replicates per time point.

Our observation of an elevated OXPHOS signature within the inner medulla is based on the transcriptional changes within the kidney tissue subjected to cold ischemia injury. While this transcriptional program suggests an increased capacity for mitochondrial respiration, gene expression alone does not necessarily reflect functional metabolic activity. It remains to be determined whether mitochondrial activity including the rate of electron transport, oxygen consumption, and ATP production as the functional output of the OXPHOS system, will mirror the transcriptional changes of the genes related to OXPHOS. Mitochondrial activity measurement techniques may require tissue dissociation which makes it challenging to implement it within intact kidney tissues. Moreover, sample preparations make it difficult to control for cold ischemia time. These technical challenges limit our ability to characterize mitochondrial flux in a compartment-specific manner. Future studies aimed at characterizing changes in mitochondrial activity during prolonged cold ischemia injury can help understand correlation between mitochondrial activity and transcriptional/proteomic changes.

Finally, there are logistical differences between the current experimental design and a clinical scenario wherein deceased donor kidneys undergo nephrectomy first and perfused *ex vivo* using kidney arteries whereas the murine kidneys in the current study were perfused through the hepatic portal vein first and harvested thereafter. Given that murine arteries present size-related challenges during perfusion, we anticipate large animal models like porcine models can be strategized to recapitulate the clinical scenario more accurately in the future. We also acknowledge that static cold storage injury likely has differences from newer storage approaches like hypothermic pulsatile perfusion and normothermic machine perfusion that may be further explored and compared in the future.

Overall, we believe this study will serve as a basis for future studies enhancing the clinical decision-making process related to selection and reduced discard of deceased donor kidneys. For example, kidney biopsies are routinely examined for underlying pathologies primarily within the cortical region. As such, pathologies developing deep within the tissue such as in the kidney inner medullary region remain poorly characterized. Our analyses suggest that this may be less of an issue in an AKI setting where our spatiotemporal ST analysis found the cortex and inner medulla to exhibit strong positive correlation, mirroring each other’s temporal molecular dynamics. However, in a cold ischemia setting where our spatiotemporal ST analysis found the cortex and inner medulla to exhibit very weak correlation, with the inner medulla displaying distinct molecular signatures potentially contributing to poor outcome, these compartmental divergences argue for targeted further investigation of inner medullary tissue to reveal potential injury mechanisms that cortex-biased assessments may miss. Furthermore, comparison of static cold ischemia vs. cold perfused vs. normothermic perfusion is also needed in the future in this evolving field. To facilitate such future comparisons as well as to make our data more accessible and explorable, we developed an interactive online browser at https://jef.works/CellCarto-ColdIschemia/. Overall, we anticipate such spatiotemporally resolved molecular comparative analysis will enhance our understanding of biological processes underlying disease pathogenesis to help elucidate novel therapeutics in the future.

## Materials and Methods

### Cold Ischemia Model and Kidney Collection

All experiments were conducted at Johns Hopkins University under an approved Institutional Animal Care and Use Committee protocol (MO24M439). 7-8 weeks old C57BL/6 male mice were anesthetized via isoflurane inhalation and perfused with ice-cold University of Wisconsin (UW) solution through the hepatic portal vein using a 10 cc syringe with a 25G needle. To facilitate circulation, the inferior vena cava below the liver was incised, enabling whole-body perfusion with UW solution. Following perfusion, the kidneys were harvested and immediately stored in UW solution at 4°C for 0, 12, 24, or 48 hours to induce cold ischemia. After the designated cold ischemia period, the kidneys were removed from the UW solution and fixed in 10% phosphate-buffered formalin for a minimum of 48 hours.

### Sample preparation for spatial transcriptomics characterization

To profile the global spatial transcriptome after cold ischemia at 0, 12, 24, and 48 hours, formalin fixed kidneys were cut sagittally, embedded in paraffin and sectioned under RNase free conditions. Kidney sections (5mm) were placed on a positively charged glass slide, H&E stained and scanned. One kidney section from each timepoint was placed on 10x Visium FFPE (formalin fixed paraffin embedded) slide using CytAssist instrument. Adjacent sections (20 mm) were used for RNA quality assessment and samples with RQN value 7 were used for spatial transcriptomic analysis. Tissue processing and downstream analyses were achieved by adopting the following 10x Genomics protocols without any alteration: (1) Visium CytAssist Spatial Gene Expression for FFPE (CG000520, Revision B) for deparaffinization, staining, decrosslinking and imaging of slides. (2) “Sequencing” sections of the Visium CytAssist Spatial Gene Expression Reagent Kits User Guide (CG000495) for hybridization. The CytAssist libraries were sequenced using a Novaseq 6000 sequencer. Gene expression counts and spot positions were obtained using the SpaceRanger pipeline (v2.0.1) for downstream analysis.

### Public data access

Previously published 10x Visium spatial transcriptomics datasets from coronal sections of female murine kidneys with bilateral ischemia reperfusion injury (ID: AKI (n=5), timepoints: sham, 4, 12, 48 hours and 6 weeks) (*20*) were obtained from the Gene Expression Omnibus (GEO) under accession GSE182939. Likewise, previously published 10x Visium spatial transcriptomics (ST) datasets sections of sagittal sections of normal control male murine kidneys (ID: CTRL(n=4)) and of male murine kidneys with bilateral ischemia reperfusion injury (ID: AKI24 (n=4), timepoint: 24 hours (all specimens)) were obtained from the authors (*22*). All datasets were accessed as gene counts matrices and spatial positions in preprocessed h5 files.

### Quality Control and Processing

Any 10x Visium spots with less than 100 counts were removed. This resulted in a total of 44,816 spots belonging to all the 17 different spatial transcriptomics datasets. R statistical software (version 4.3.1 (2023-06-16)) was used for this purpose and all the computational analyses hereafter.

### Data Integration, Batch Correction and Graph Based Clustering

For data integration, for each of the individual ST dataset (prior to CPM normalization), we removed genes with less than 10 reads across all spots. Thereafter, we feature selected for over-dispersed genes in each dataset using the getOverdispersedGenes() function from MERINGUE (*32*) To mitigate sample-specific effects, we identified genes that were overdispersed in at least 2 datasets resulting in 1,378 final shared overdispsersed genes. Using these over-dispersed genes, the top 30 principal components (PCs) were identified. Batch correction was performed on the PCs to obtain harmonized PCs using the HarmonyMatrix() function (parameters: theta = 10, lambda = 0.05) from Harmony (*23*). Low-dimensional 2D embeddings of the initial and harmonized PCs were obtained using tSNE (*33*). Graph-based Louvain clustering was performed on the harmonized PCs to identify transcriptionally similar spots using getClusters() function (parameters: k=100) from MERINGUE (*32*). Thereafter, for normalization purposes and subsequent linear regression analysis (covered in next section), the gene list was restricted to 19,454 unique genes that were shared across all 17 ST datasets.

### Linear regression modeling

Gene count matrices of CIS tissue specimens containing 19,454 unique shared genes (rows) at each timepoint 0, 12, 24 and 48 hours were subset into smaller compartment-specific matrices based on the annotations obtained from harmonized clustering. This resulted in generation of 3 matrices per timepoint capturing all the spots within the inner medulla, outer medulla, and cortex compartments respectively. Thereafter, we performed CPM (counts per million) normalization on the gene counts matrices to obtain count-normalized matrices using normalizeCounts() function from MERINGUE (*32*). To avoid over-representation from any particular compartment (especially the cortex), 190 random spots were sampled for each of these matrices belonging to the 3 compartments (per timepoint) to perform compartment-specific linear regression at each timepoint. lmfit() function from limma (*34*) package was used to perform linear regression with time (encoded as log_e_(hours+1)) as the independent variable (x) and count-normalized gene counts as the dependent variable (y). Goodness of fit value, R^2^ (R^2^>0.026) and adjusted p-value (adj. p< 0.05) was used as a criterion to select the genes (CIS) whose expression had significant temporal association (reflected by the magnitude of linear regression slope with unit as: CPM normalized read/log_e_(hours+1)). Similar procedure was implemented on AKI tissue specimens at timepoints (0 (sham), 4, 12 and 48 hours) with R^2^>0.093 and adj.p<0.05. The choice of R^2^ values for CIS and AKI tissue specimens was based on a bootstrap sampling experiment, where we randomly selected 190 spots from each compartment belonging to the CIS or AKI tissue and performed linear regression on those spots (CPM normalized gene expression vs log_e_(time+1) (cold ischemia time or warm ischemia time)). We permutated this process for 100 times and collected the R^2^ and p-value for each gene for all the 100 permutations. Thereafter, we ranked all the gene based on their consistency score (i.e., Number of times the p-value for the gene was less than 0.05 out of 100 times). Then we created a cut-off at the 95th quantile of this ordered gene list and the mean R^2^ of the gene (average R^2^ for the gene from all the 100 permutations) at the 95^th^ quantile mark was identified as the R^2^ cut-off for qualifying genes as genes with significant temporal regulation within a compartment, identified through the linear regression analysis. Finally, the largest R^2^ cut-off value out of the three compartments was selected as the R^2^ cut-off for either CIS (i.e, 0.026) or AKI (i.e., 0.093) tissue specimens.

### Gene Set Enrichment Analysis

These genes with significant temporal association identified through linear regression were ordered based on the strength of their temporal association (linear regression slopes). These genes were assigned a value which reflected their rank in the ordered list such that temporally upregulated genes received a positive score while temporally downregulated genes received a negative score. Thereafter, gene set enrichment analysis was performed on the ordered list of genes along with their assigned values using gseKEGG() function from ClusterProfiler (*35*) package and GSEA() function from Msigdbr (*36*) package to identify enriched KEGG and Hallmark (gene set of Msigdbr) molecular pathways respectively. Adjusted p-value (adj. p<0.05) was used as a cut-off criterion to select significantly enriched molecular pathways. Running enrichment score plots were plotted using gseaplot2() functions from enrichplot (*37*) package. Benjamini-Hochberg method was used for multiple hypothesis testing for gene set enrichment analysis.

### Compartmental correlation analysis and linear regression

Correlation analysis of the temporal transcriptomic dynamics between the different compartments was performed using cor.test() function in R to obtain Pearson correlation coefficient (PCC) and p-value. Linear regression was performed using lm() function in R and summary function was used to obtain the R^2^ value.

### Differential Expression Analysis

Differential expression analysis was performed to identify top marker genes of inner medulla, outer medulla, and cortex compartments. Briefly, count matrices (raw counts) belonging to all the 17 ST datasets were split into three groups based on the labels (from earlier assigned compartmental annotations) of the spots. Thereafter, differential expression analysis was performed between the interest group and the other two groups pooled together into one group using DESeq2 (*38*). Top 30 genes based on their log2 fold change (and adjusted p-value<0.05) from each of the compartments were identified and plotted as heatmaps. Differential expression was also performed between the inner medulla compartment of all 4 kidney tissue specimens belonging to normal control kidneys (CTRL) and 4 warm ischemia reperfusion injury (AKI24) to evaluate consistency with linear regression modeling.

### Cell type Deconvolution

Reference-free cell type deconvolution of 10x Visium spots of the cold ischemia kidney tissue was performed using STdeconvolve (*39*). Gene count matrices of all cold ischemic kidney tissue across timepoints were collated to jointly deconvolve all time points. Feature selection was limited to previously identified shared overdispersed genes. We set K=7 to deconvolve 7 major cell types within the kidney tissue. The choice of 7 cell types was taken based on findings from previous work (*20*). Deconvolved cell-type proportions were visualized using scatterbar (*40*). Cell-type-specific marker genes identified for the respective deconvolved cell types and ranked by their normalized expression levels (expression level of a particular gene was normalized across all deconvolved cell types) were used for gene set enrichment analysis using gseGO() function from clusterProfiler package.

### GC content and 3’ UTR Length Calculations

The GC content and 3’ UTR length for all the different transcripts belonging to all the genes were derived using BiomaRt package (*41*, *42*). Thereafter, the GC content and 3’ UTR length for all the transcripts belonging to a particular gene were averaged respectively to obtain the averaged gene level GC content and 3’ UTR length values. These averaged gene level GC content and 3’ UTR length have been used for comparative analysis between OXPHOS genes and all other temporally regulated genes within the inner medulla of cold ischemia kidney tissue.

### RNA isolation and qPCR

Kidney cortex and medulla (inner + outer medulla) were collected as previously described (*43*). Total RNA was isolated from samples using a QIAshredder (Qiagen, Valencia, CA, Cat.No.79654) and a RNeasy Mini kit (Qiagen, Cat.No.74104) and reverse transcribed using High-Capacity cDNA Reverse Transcription Kit (Applied Biosystems, Foster City, CA, Cat.No. 4368814). Real-time PCR was performed in QuantStudio 12K Flex (Applied Biosystems, Foster City, CA) using the SYBR Green PCR master mix (Applied biosystems, Cat.No. 4309155). *β*-actin served as the reference gene and the relative mRNA fold change expression values were calculated using a DD cycle threshold method. Two-sided Wilcoxon rank sum test was used to compare the relative mRNA fold change values for *Uqcr11*, *Cox6a1* and *Pink1* between the cortex and medulla regions of the kidney using wilcox.test() function. The primer sequences for genes are listed below:

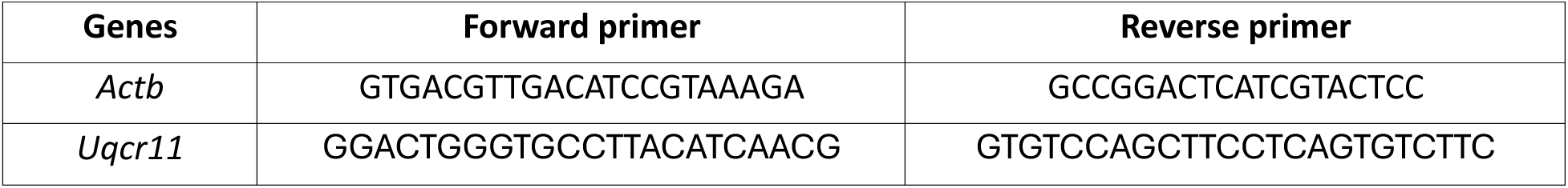
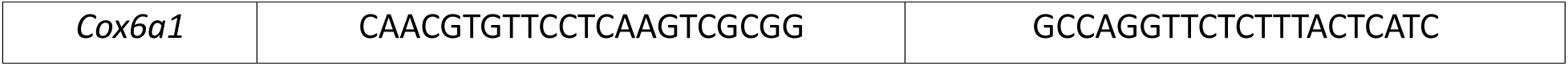

### Immunostaining and Immufluorescenece microsocopy

The paraffin embedded sections were deparaffinized and hydrated by incubation in xylene and subsequently in 100%, 95%, 80%, 70% ethanol and deionized water. Thereafter, antigen retrieval was performed by microwaving the deparaffinized specimens in 1x Tris EDTA buffer (ThermoFisher Scientific, Catalog no: 00-4956-58) for a minute. The specimens were permeabilized using 0.1% (v/v) Triton X-100 in phosphate buffered saline (PBS) for 5 minutes. Primary antibodies used for different protein markers were: Uqcr11 (ThermoFisher Scientific, Catalog no: 14793-1-AP), Pkm (Abcam, Catalog no: AB154816), Slc2a1 (Abcam, Catalog no: AB150299). Primary antibody solution was prepared in 0.1% (v/v) Tween-20 in PBS containing 10% (v/v) normal donkey serum (Sigma Aldrich, D9663). All primary antibodies (host species: Rabbit) were used at a dilution of 1:100 (v/v). Goat Anti-Rabbit IgG Alexa Fluor 647 (Abcam, Catalog no: AB150083) was used as a secondary antibody and was used for conjugation with all primary antibodies used in this study at a dilution of 1:200 (v/v). The immunostained slides were imaged using Nikon Suno Spinning disk microscope and all images were analyzed using Fiji.

## Code availability

Code to reproduce the analyses and results of this study is available on GitHub at: https://github.com/ssingh95jhu/Cold_Ischemia_Injury_Molecular_Characterization_v2

An interactive browser is available at https://jef.works/CellCarto-ColdIschemia/ with source code at https://github.com/JEFworks-Lab/CellCarto-ColdIschemia

## Data availability

Count matrices, spatial positions, and H&E images for the cold ischemia Visium datasets generated in this study have been deposited in a Zenodo repository at https://doi.org/10.5281/zenodo.15359609

## Acknowledgements

This material is based upon work supported by the National Institute of General Medical Sciences of the National Institutes of Health under Award Number R35-GM142889 (SS, JF) and the National Science Foundation under Grant No. 2047611 (DV, JF). H.R. and S.N. were supported by NIH/NIDDK grants (R01DK132278, R01DK123342, U54DK137331) and the Kidney precision medicine project biomarker award. We thank Dr. Ian Dobbie and Yuan Cai at the Integrated Imaging Center (IIC), Johns Hopkins University for their insights and feedback related to microscopic imaging and analysis of immunostained slides.

## Contributions

J.F. and H.R. conceptualized the study. S.S. led the data analysis, interpretation, and writing under the guidance of J.F. and with input from S.K.P., R.M., S.N., and H.R. S.K.P., R.M. and S.N. performed the cold ischemia protocol, kidney tissue collection, sectioning and tissue processing for spatial transcriptomics under the guidance of Z.S. and H.R. S.K.P., R.M. and S.N. led the qPCR validation under the guidance of H.R. S.S. led the immunostaining validation. D.V. developed the web application under the guidance of S.S. and J.F. All authors contributed to the writing of the manuscript. All authors approved the final manuscript.

## Supplemental Figures

**Figure S1.**
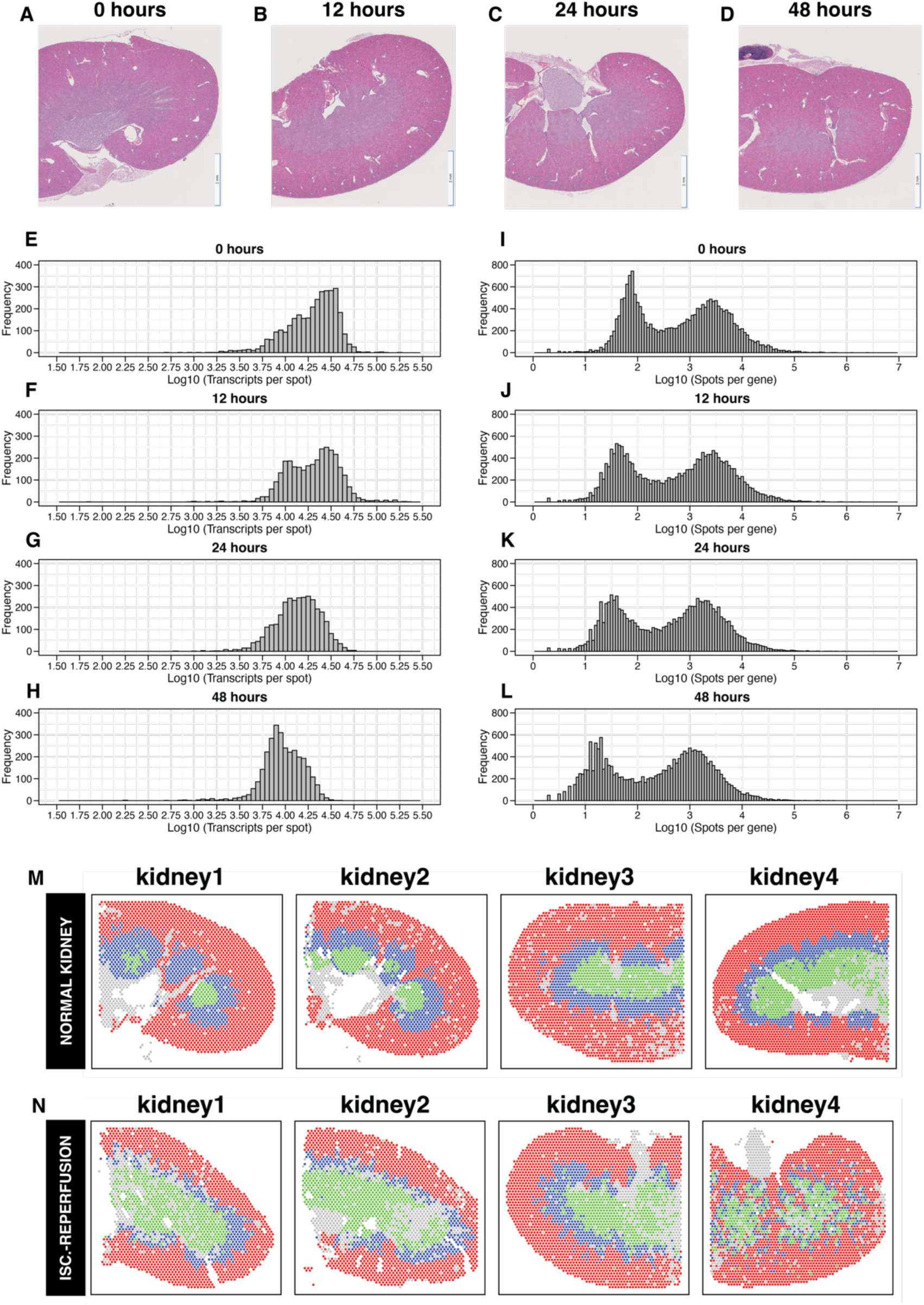
Tissue specimens and quality control of cold ischemia injury kidneys. H&E-stained images of kidney tissue specimens subjected to **(A)** 0 hours **(B)** 12 hours **(C)** 24 hours and **(D)** 48 hours of cold ischemia injury. Histograms depicting distribution of **(E-H)** transcripts per spot and **(I-L)** spots per gene at time points 0 hours, 12 hours, 24 hours and 48 hours of cold ischemia injury respectively. Annotations highlighting the different compartments namely cortex (red), outer medulla (blue), inner medulla (green) and other (gray), within the normal control (CTRL) kidneys (**M**) and 24-hours warm ischemia reperfusion (AKI24) injury (**N**) tissue specimens.

**Figure S2.**
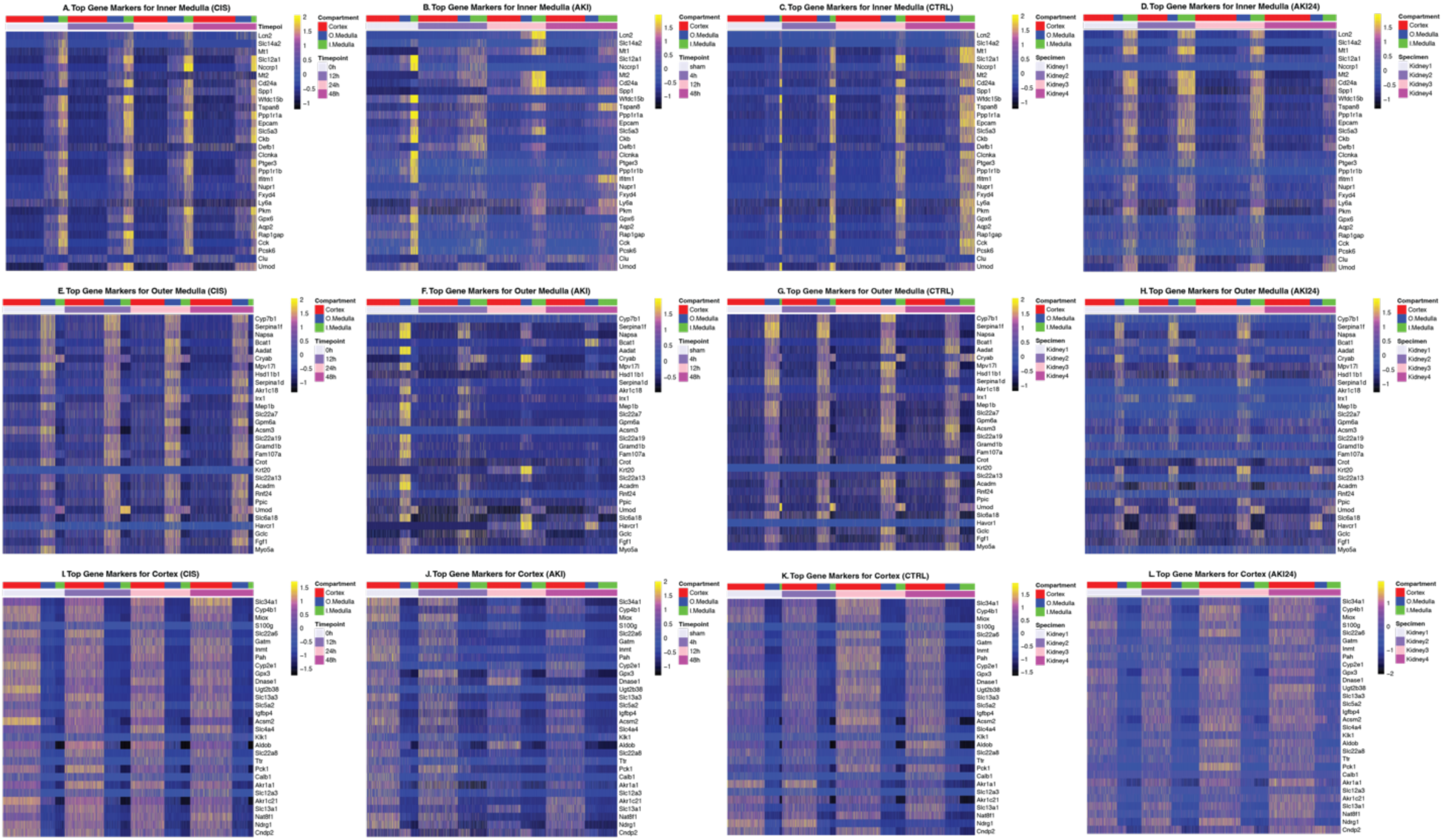
Heatmap of top compartment-specific marker genes for cold ischemia (CIS), warm ischemia-reperfusion-induced acute kidney injury (AKI), normal control kidneys (CTRL) and 24-hours warm ischemia-reperfusion-induced acute kidney injury (AKI24) injury datasets. Heatmap of top compartment-specific marker genes expression within the different kidney tissue compartments namely **(A,B,C,D)** inner medulla (I.medulla (green)), **(E,F,G,H)** outer medulla (O. medulla (blue)) and **(I,J,K,L)** cortex (red) belonging to different timepoints of cold ischemia (CIS) and warm ischemia-reperfusion (AKI) injury, normal control kidneys (CTRL) and 24-hours warm ischemic-reperfusion injury (AKI24) datasets respectively.

**Figure S3.**
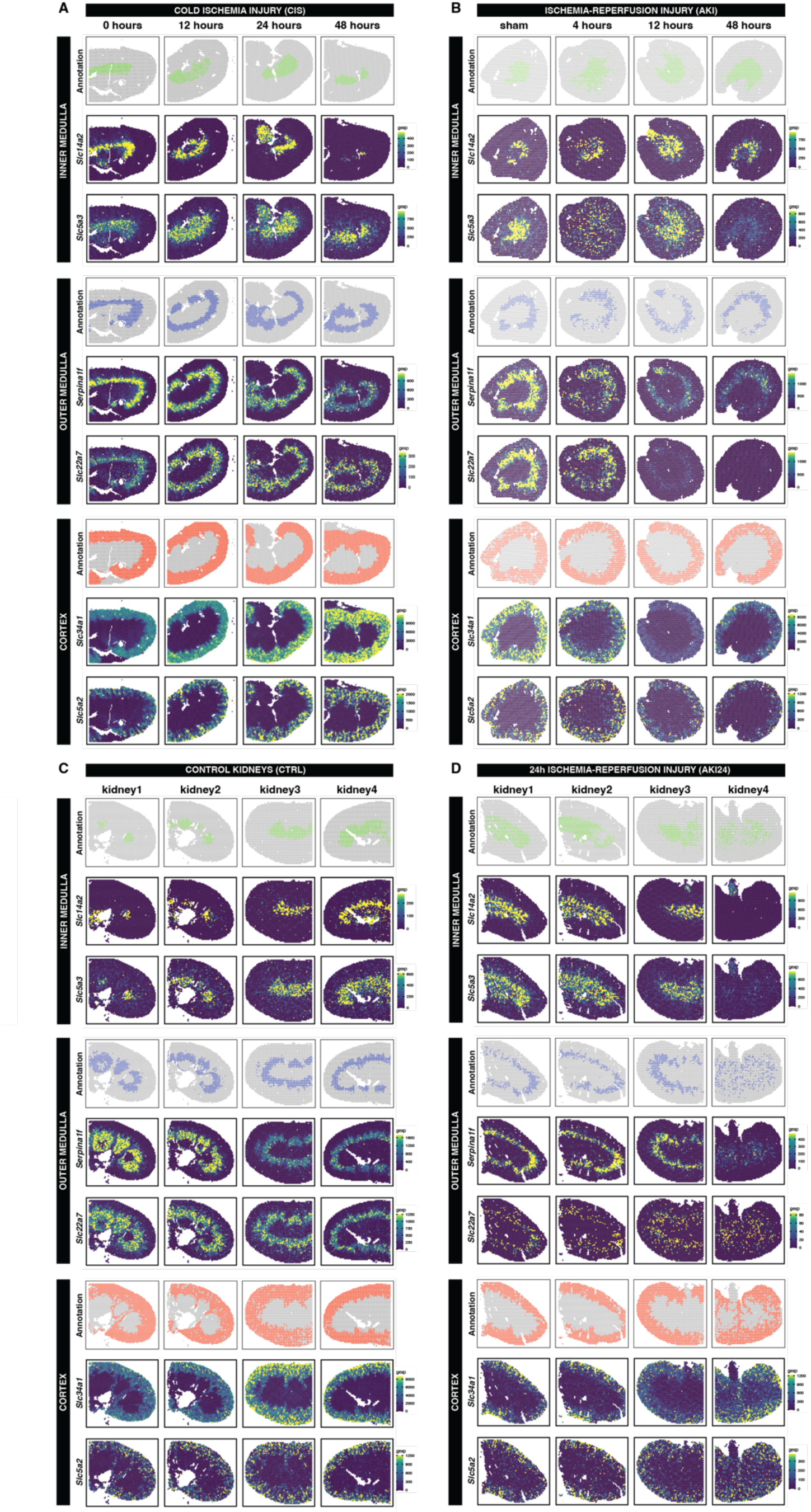
Spatiotemporal expression patterns of compartment-specific marker genes in different experimental groups. Presentation of inner medulla marker genes (*Slc14a2*, *Slc5a3*); outer medulla marker genes (*Serpina1f, Slc22a7*); cortex marker genes (*Slc34a1, Slc5a2*) within the **(A)** cold ischemia injury (CIS), **(B)** warm ischemia-reperfusion injury (AKI), **(C)** normal control kidneys (CTRL) and **(D)** 24-hours warm ischemia-reperfusion injury (AKI24) kidney tissue specimens. Color coded annotations: red (cortex), blue (outer medulla), green (inner medulla) and other (gray), highlight the location of the different compartments within respective tissue specimens.

**Figure S4.**
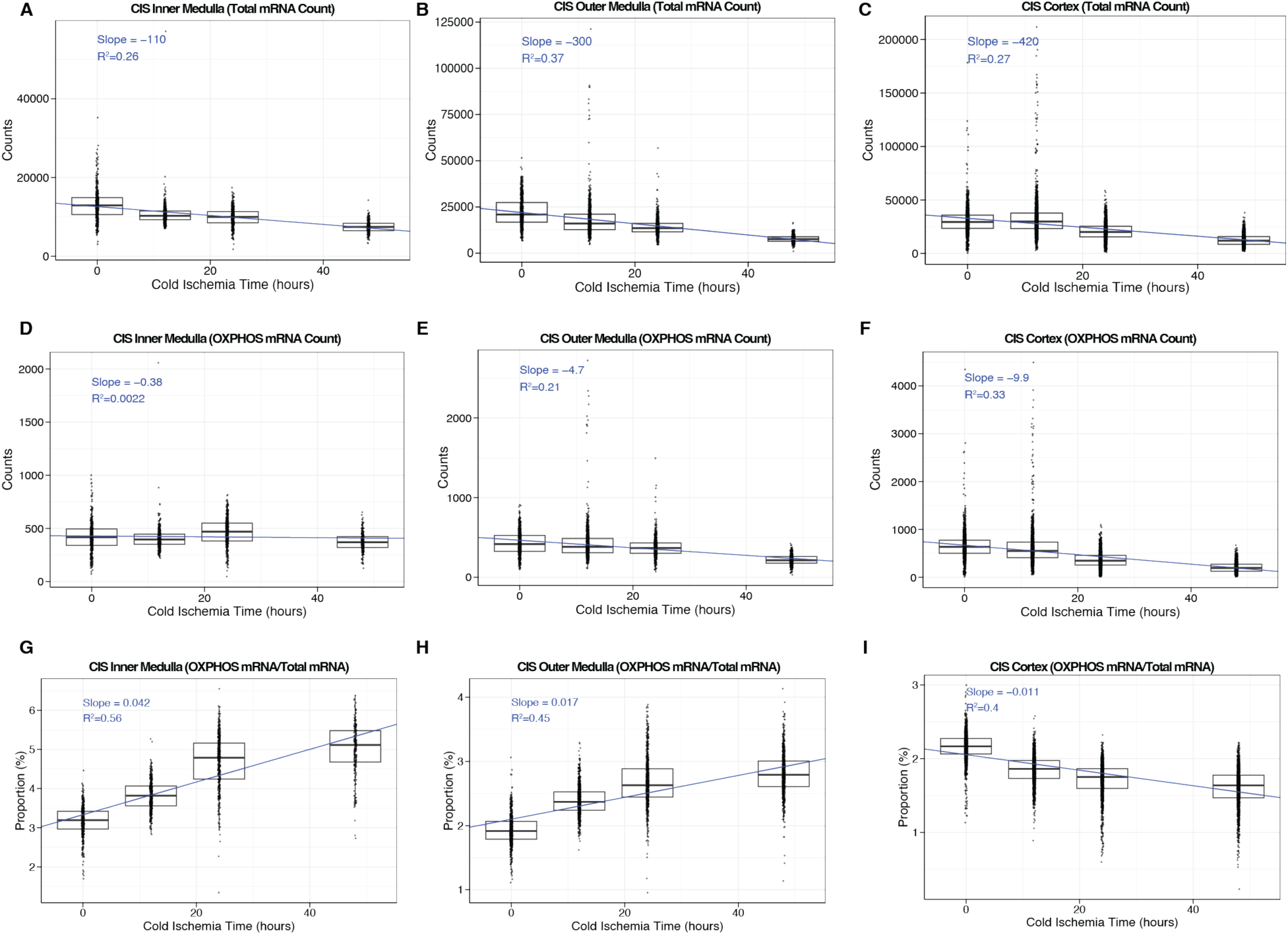
Compartment-specific proportional enrichment or depletion of OXPHOS genes within cold ischemia kidney tissue. Temporal change in total mRNA count within the **(A)** inner medulla **(B)** outer medulla and **(C)** cortex. Temporal change in OXPHOS related genes within the **(D)** inner medulla **(E)** outer medulla and **(F)** cortex. Temporal change in proportion of OXPHOS related genes with **(G)** inner medulla **(H)** outer medulla and **(I)** cortex. The regression slopes (Slope) and goodness of fit (R^2^) of each of the plots is provided at the upper left corner of the respective plots.

**Figure S5.**
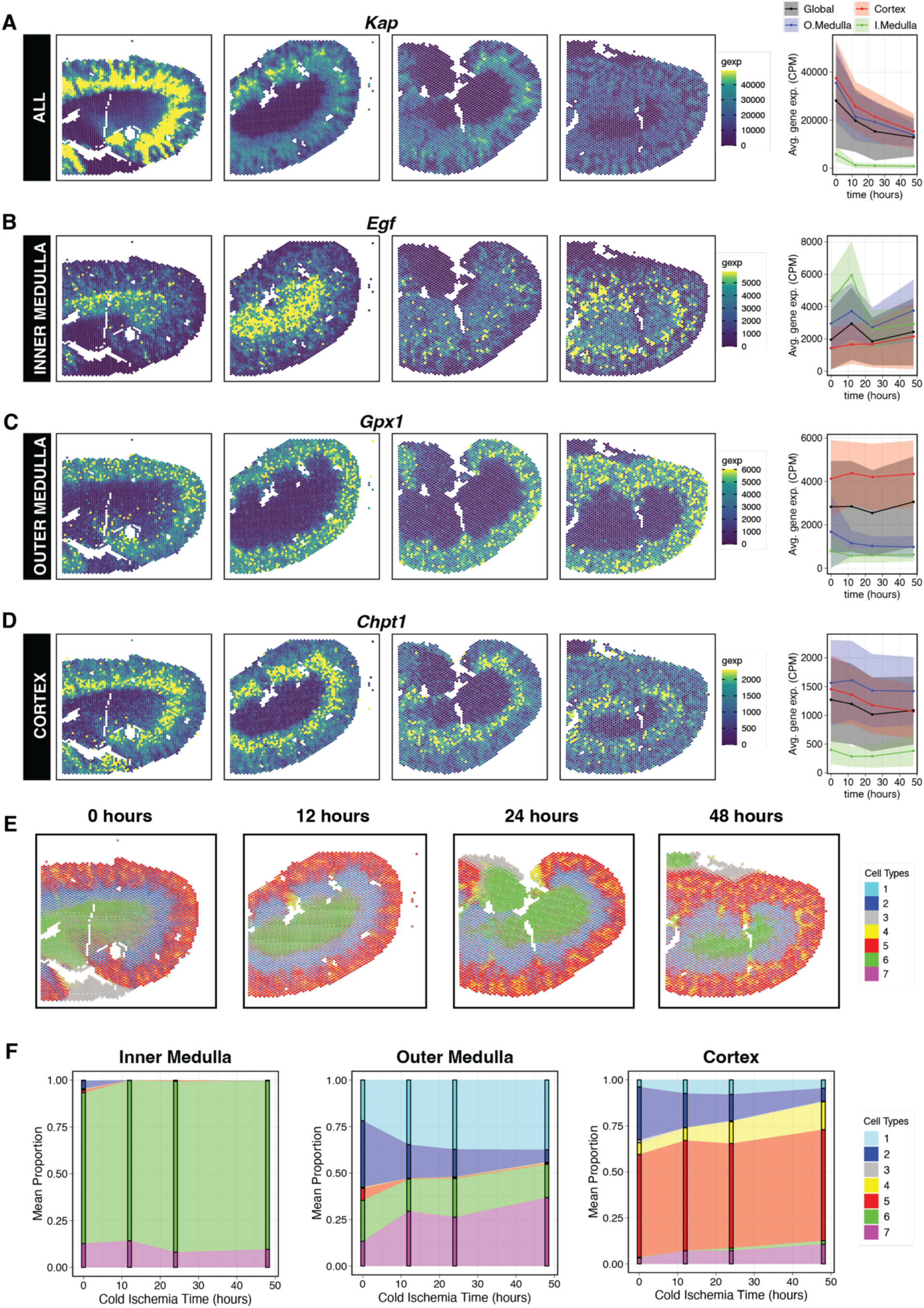
Representative genes demonstrating compartment-specific downward temporal trend in cold ischemic kidneys and deconvolved cell type changes. While (A) *Kap* demonstrated temporally downward trend in all three compartments with varying degrees, (B) *Egf,* (C) *Gpx1* and (D) *Chpt1* were confined to the inner medulla, outer medulla, and cortex respectively. The ribbon plots demonstrate the mean ± s.d. expression (CPM normalized) of the respective genes within different compartments of the corresponding cold ischemic tissue at different time points. **(E)** Spatial gene expression plots demonstrating spatiotemporal expression of deconvolved cell types (1–7) within cold ischemic kidney tissues. **(F)** Compartment-specific deconvolved cell type composition changes during cold ischemia injury in kidneys.

**Figure S6.**
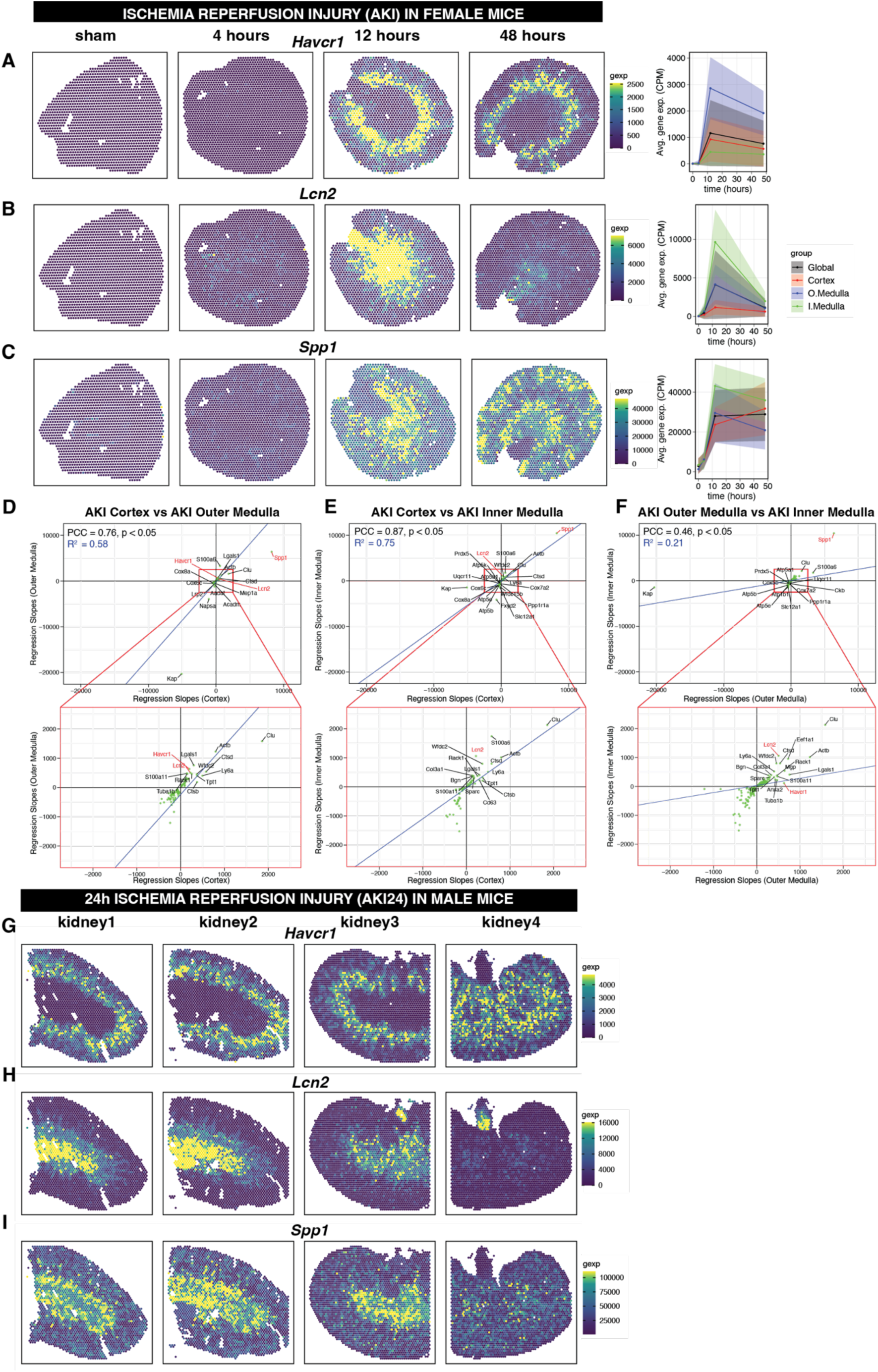
Spatiotemporal transcriptional dynamics within female and male experimental models of warm ischemia-reperfusion injury (i.e., AKI and AKI24). Spatial plots depicting the spatiotemporal expression of putative AKI marker genes like (A) *Havcr1*, (B) *Lcn2* and (C) *Spp1* within warm ischemia-reperfusion injury (AKI) kidney tissue specimens from female murine models. Scatter plots depicting the correlation analysis of the temporal transcriptional dynamics between the different compartments: **(D)** cortex vs outer medulla **(E)** cortex vs inner medulla and **(F)** outer medulla vs inner medulla of the acute kidney injury (AKI) tissue. Genes exhibiting covarying (green) or divergent (purple) temporal trend within these compartments can be visualized along with their Pearson correlation coefficients (PCC) and linear regression slopes (blue line). AKI marker genes like *Spp1*, *Havcr1* and *Lcn2* within these compartments have been highlighted red. Spatial plots depicting the spatiotemporal expression of (G) *Havcr1*, (H) *Lcn2* and (I) *Spp1* within 24-hours warm ischemia-reperfusion injury (AKI24) kidney tissue specimens from male murine models.

**Figure S7.**
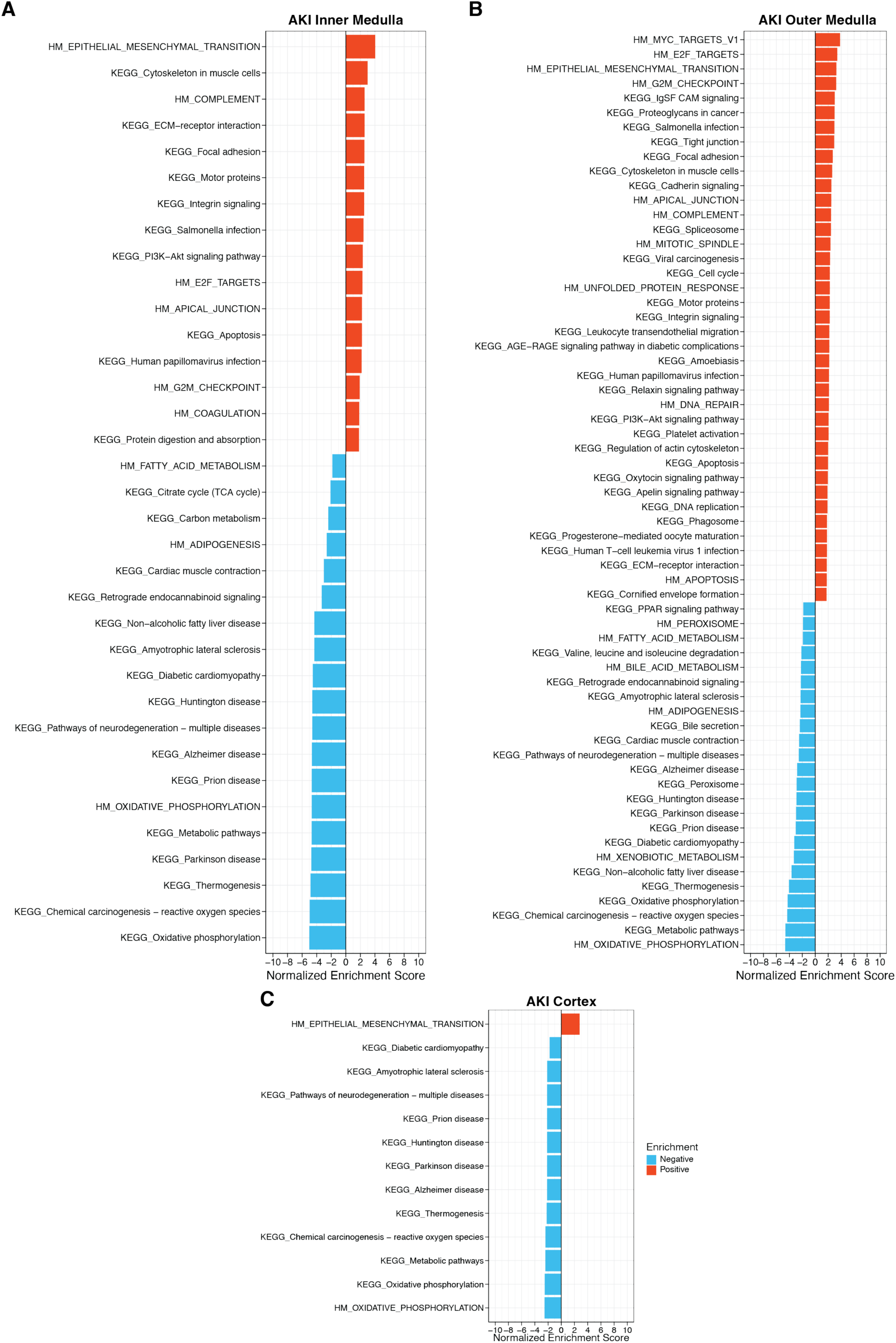
Compartment-specific enrichment of molecular pathways within acute kidney injury (AKI) tissues. Bar plots highlighting all the enriched molecular pathways (Hallmark and KEGG) that demonstrated similar temporal dynamics (enrichment: positive (red), negative (cyan)) within the corresponding **(A)** inner medulla, **(B)** outer medulla and **(C)** cortex of warm ischemia-reperfusion injury (AKI) kidney tissue. Similar, bar plots highlighting all the enriched molecular pathways (KEGG) that demonstrated similar temporal dynamics (enrichment: positive (red), negative (cyan)) within the corresponding **(D)** inner medulla, **(E)** outer medulla and **(F)** cortex of warm ischemia-reperfusion injury (AKI) kidney tissue. (Note: Hallmark pathways have a prefix “HM_” and are in uppercase whereas KEGG pathways have a prefix “KEGG_”.)

**Figure S8.**
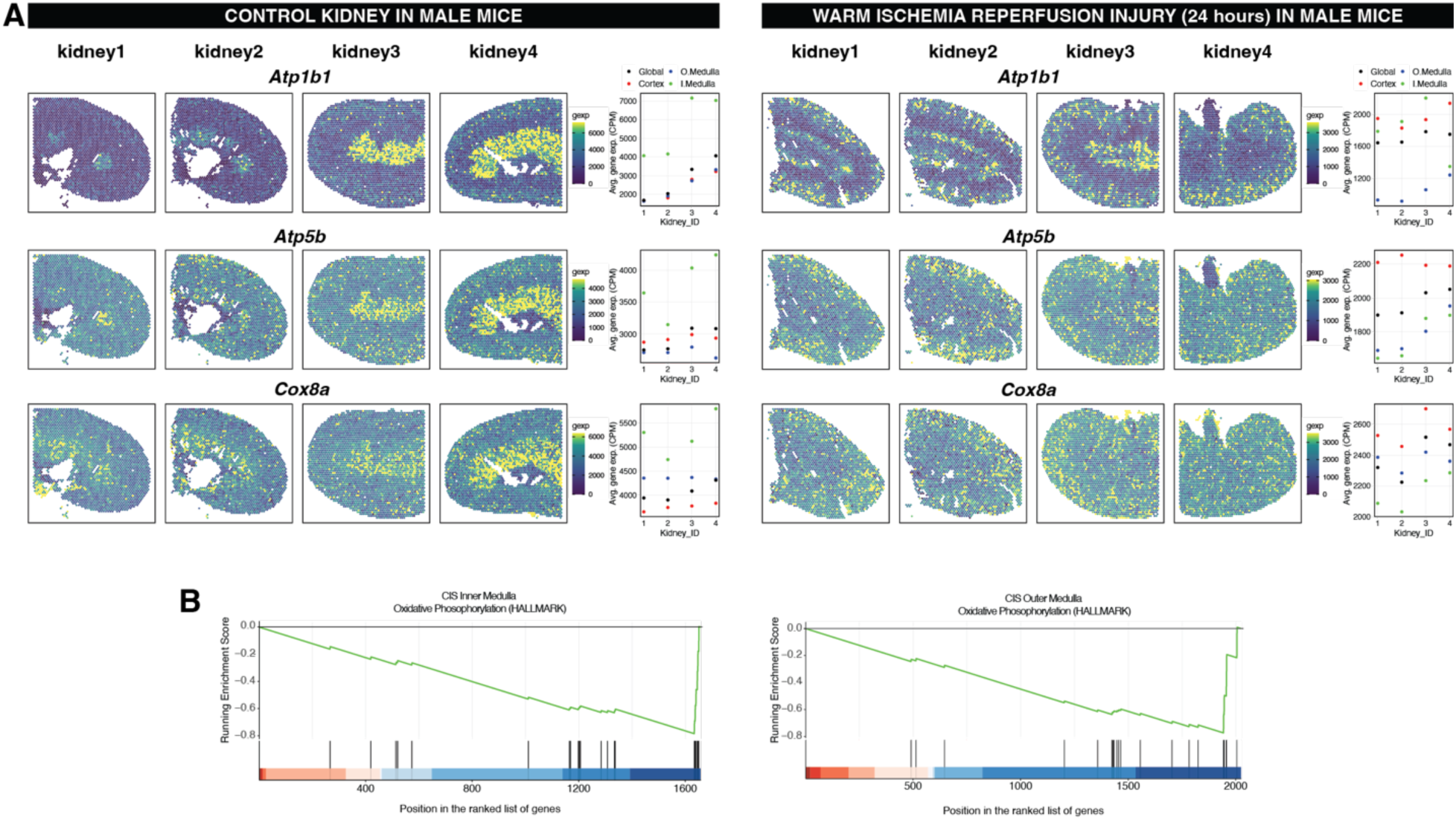
Differential spatiotemporal expression of OXPHOS genes between the control (CTRL) and warm ischemia injury (24 hours) male kidney. **(A)** Spatial plots depicting the differential spatiotemporal expression of OXPHOS genes like *Atp1b1, Atp5b* and *Cox8a* between the control and warm ischemia-reperfusion injury (AKI24) kidney tissue specimens from male murine model. **(B)** Running enrichment score plots highlighting differential negative enrichment of OXPHOS pathway (HALLMARK) within the inner medulla of warm ischemia-reperfusion injury (AKI24) male kidney tissue as compared to control male kidney.

